# An allosteric switch between the activation loop and a c-terminal palindromic phospho-motif controls c-Src function

**DOI:** 10.1101/2022.10.16.512342

**Authors:** Nicolás Cuesta, Julia Contreras, Jana Sánchez-Waldermer, Pablo Soriano-Maldonado, Ana Martín-Hurtado, Inés G. Muñoz, Marta Llimargas, Javier Muñoz, Iván Plaza Menacho

**Affiliations:** Protein Phosphorylation and Cancer Group, Structural Biology Programme, Spanish National Cancer Research Center (CNIO), C/Melchor Fernández Almagro num. 3, 28029 Madrid, Spain; Proteomics Unit, Spanish National Cancer Research Center (CNIO), C/Melchor Fernández Almagro num. 3, 28029 Madrid, Spain; Protein Crystallography Unit, Spanish National Cancer Research Center (CNIO), C/Melchor Fernández Almagro num. 3, 28029 Madrid, Spain; Institute of Molecular Biology of Barcelona; Department of Structure of Macromolecules. National Biotechnology Centre (CNB-CSIC), C/Darwin 3, 28049, Madrid, Spain; IIS Biocruces Bizkaia. Building Biocruces Bizkaia 1, 48903 Cruces, Bizkaia. Ikerbasque, Basque Foundation for Science

## Abstract

Auto-phosphorylation controls the transition between discrete functional and conformational states in protein kinases, yet the structural and molecular determinants underlaying this fundamental process remain unclear. Here we show that c-terminal Tyr 530 is a *de facto* c-Src auto-phosphorylation site with slow time-resolution kinetics and strong intermolecular component. On the contrary, activation-loop Tyr 419 undergoes fast kinetics and a cis-to-trans phosphorylation-switch that controls c-terminal Tyr 530 auto-phosphorylation, enzyme specificity and strikingly, c-Src non-catalytic function as a substrate. In line with this, we visualize by X-ray crystallography a snapshot of Tyr 530 intermolecular phosphorylation in which a c-terminal palindromic phospho-motif flanking Tyr 530 on the susbtrate molecule engages the P-loop of the active kinase for ready entry prior catalysis. Perturbation of the phospho-motif accounts for c-Src disfunction as indicated by viral and a colorectal cancer (CRC) associated c-terminal deleted variants. We show that c-terminal residues 531 to 536 are required for c-Src Tyr 530 and global auto-phosphorylation, and this detrimental effect is caused by the susbtrate molecule inhibiting allosterically the active kinase. Our work reveals a bi-directional crosstalk between the activation and c-terminal segments that controls the allosteric interplay between susbtrate and enzyme acting kinases during auto-phosphorylation

**Highlights:**

1. A bi-directional phospho-switch connecting the activation and c-terminal segments controls c-Src function
2. Activation-loop Tyr 419 is required for c-terminal Tyr 530 auto-phosphorylation, enzyme specificity and non-catalytic function as a substrate
3. Biochemical and structural visualization of c-Src Tyr 530 intermolecular auto-phosphorylation
4. A double-phosphorylated c-Src on both the activation and c-terminal segments is a fully active protein
5. Cancer associated c-terminal deleted variants inhibit allosterically c-Src activity by a dominant negative effect

## INTRODUCTION

In 1980, Hunter and Sefton found that the products of the Rous Sarcoma Virus transforming gene v-Src and its cellular homolog c-Src phosphorylated tyrosine residues on target proteins (Hunter, 2015; Hunter and Sefton, 1980). This was the first report of a protein kinase with tyrosine specificity, a seminal work for the cancer research field that proceeded the discovery of many other viral oncogenes that encoded proteins with tyrosine kinase activity (Lipsick, 2021). The further identification of oncogenic mutations in human genes that codified receptor tyrosine kinases acting as drivers in human cancers such us lung (EGFR), breast (HER2), gastric (c-Kit), renal and hepatocellular (MET) and thyroid (RET) cancers among many others, illustrated the causative role of this family of proteins in cancer (Blume-Jensen and Hunter, 2001; Torkamani et al., 2009). Over the last thirty years, the structural and molecular understanding of protein kinases has allowed the design and development of an important number of new anti-cancer drugs including small-molecule inhibitors and monoclonal antibodies that revolutionized targeted and personalized therapies in oncology e.g. Gleevec and Herceptin (Druker et al., 1996; Slamon et al., 2001)

c-Src belongs to the Src family of kinases (SFKs) composed of eleven structurally similar non-receptor protein tyrosine kinases: Src, Fyn, Lyn, Yes, Blk, Lck, Hck, Fgr, Srm, Brk and Yrk (Manning et al., 2002), all of which share a conserved arrangement of four different functional domains, named Src homology (SH) domains as well as a regulatory c-terminal sequence. The intrinsically disordered N-terminal region SH4 domain undergoes myristylation on a conserved glycine required for membrane attachment and localization, regulation of kinase activity and intracellular stability (Buser et al., 1994; Resh, 1994). SH3 and SH2 domains allow inter-as well in intra-molecular interactions with proline-rich (Feng et al., 1994; Musacchio et al., 1994; Yu et al., 1994) and phospho-tyrosine containing sequences (Waksman et al., 1992; Waksman et al., 1993) respectively that are critical for c-Src functional regulation. The SH1 catalytic domain display intrinsic tyrosine kinase activity, which is further regulated by a c-terminal regulatory sequence (Xu et al., 1997). c-Src activity is regulated by phosphorylation and protein-protein interactions events. In the current paradigm, phosphorylation at Tyr 530 by c-terminal Src kinase (CSK) is inhibitory, while phosphorylation on Tyr 419 in the activation loop is activating, although neither of these phosphorylation sites by themselves exert full positive or negative regulatory control (Roskoski, 2004, 2005). In a cellular context, dimerization appears to substantially enhance auto-phosphorylation and the phosphorylation of selected substrates, while interfering the dimerization precludes catalytic function (Spassov et al., 2018). Structural studies have revealed that c-Src is held in an autoinhibited monomeric state by intramolecular interactions between the SH2 domain and the CSK-phosphorylated Tyr 530 on the c-terminal segment, while the SH3 domain engages a proline-rich linker between the SH2 and kinase domain N-lobe where the active site is disrupted by displacement of the αC helix (Xu et al., 1997). These intramolecular connections can be displaced by intermolecular interactions between the SH2 and SH3 domains with higher affinity ligands (Moarefi et al., 1997) or by de-phosphorylation of Tyr 530 by a number of phosphatases (Bjorge et al., 2000; Somani et al., 1997). Auto-phosphorylation on Tyr 419 stabilizes the activation loop into an active conformation favorable for substrate binding (Porter et al., 2000; Schindler et al., 1999) and intermolecular phosphorylation (Boerner et al., 1996; Cooper and MacAuley, 1988; Meng and Roux, 2014). However, the precise role of Tyr 419 auto-phosphorylation in catalytic function and signalling is paradoxically not yet fully understood. In fact, the precise mechanism by which auto-phosphorylation regulates c-Src intrinsic activity and conformational state independent of external inputs, and how this process is corrupted in cancer remains elusive.

SFKs expression and activity is upregulated in solid tumors and some hematologic malignancies, often correlating with progressive stages of the diseases (Kim et al., 2009). However, c-Src activating mutations or amplifications are rare in human cancers (Bailey et al., 2018; Curtis et al., 2012), with the exception of colorectal cancer (Irby et al., 1999). Constitutive c-Src activation in human cancers cells appears to be driven at least partially by secondary alterations in upstream activators. c-Src is activated by a plethora of receptor tyrosine kinases (RTKs) including epidermal growth factor receptor (EGFR), fibroblast growth factor receptor (FGFR) and insulin-like growth factor-1 receptor (IGF1R), among others (Parsons and Parsons, 1997). Gain of function mutations and/or overexpression of these RTKs cooperate and synergize with c-Src activation to promote tumorigenesis probably by releasing c-Src from a closed autoinhibited state (Bromann et al., 2004). On the other hand, CSK mediated phosphorylation of c-terminal Tyr 530 inactivates c-Src only when the protein is not previously auto-phosphorylated (Sun et al., 1998), which indicates the existence of intrinsically coordinated and sequential auto-phosphorylation events coupled to different conformation and functional states. These data also suggest that perturbation of auto-phosphorylation could account for the paradoxically high levels of dually-phosphorylated c-Src found in malignant cells (Sun et al., 1998). Alternatively, c-Src could function independently of CSK and the latter might play other roles rather than regulating the former (see discussion). Here we provide functional and structural evidence related to the identification of new *de facto* c-Src auto-phosphorylation sites and how “canonical” activating and repressive tyrosine residues are actually playing other important roles and functions not previously envisioned. Our data dissect a sequential and coordinated cis-to-trans phosphorylation switch connecting the activation and c-terminal segments that controls c-Src enzymatic specificity and non-catalytic functions as a susbtrate. Evaluation of cancer related c-terminal variants provide evidence of the existence of a complex allosteric node at the c-terminus of c-Src controlling the crosstalk between susbtrate- and enzyme-acting kinases, and how perturbation of this allosteric phospho-switch drive c-Src dysfunction in cancer by a dominant negative effect.

## MATERIALS AND METHODS

### Plasmids

A pET28a-TEV plasmid codifying an N-Terminal 6xHis tag followed by a Tobacco etch virus (TEV) protease recognition site was used to express human c-Src (UniProtKB P12931) three domain (3D, SH3-SH2-KD aa 84-536) or kinase domain (KD, aa 254-536) codifying sequences. Both pFCDUET-YopH phosphatase and pGKJE8-GroEl/GroEs chaperones plasmids (Takara chaperone plasmid set cat. # 3340) were used for co-expression experiments. These constructs were a kind gift of Dr. Daniel Lietha (CIB, Madrid). For expression of recombinant drosophila c-Src isoform 42A (Src42A, UniProtKB Q9V9J3) we used a pET-His10-Src42A plasmid (Addgene #126674) codifying a full-length drosophila isoform (aa 1-517) with a N-terminal 10x His tag and a Thrombin protease recognition site. For experiments in Drosophila a pUAST-attB plasmid encoding drosophila Src homolog isoform A (Src42A) was cloned by amplifying Src42A codifying sequence from a pGEX-Src42A donor plasmid (Addgene #126673) using primers with EcoRI (forward 5′-CTGAATAGGGAATTGGGAATTCATGGGTAACTGCCTCACC-3′) and XbaI (reverse 5′-CCTTCACAAAGATCCTCTAGATCAGTAGGCCTGCGCCTC-3′) restriction sites.

### Site-directed mutagenesis

Site-directed mutagenesis was performed in order to generate all the point mutants and variants described in this study using the Q5-site directed mutagenesis kit (New England Biolabs) following manufacturer instructions and the indicated primers (see extended material and methods).

### Expression and purification of recombinant proteins

In order to express high yields of soluble and monodisperse recombinant human c-Src protein we followed a modified protocol, in which in addition to YopH phosphatase (Seeliger et al., 2005), c-Src was also co-expressed with the chaperone GroEL. For co-expression, pET28a-TEV-hSrc [84-536] and pET28a-TEV-hSrc [254-536] plasmids were co-transformed in E. coli BL21 bacteria strand previously transformed with pFCDUET-YopH and pGKJE8-GroEl/GroEs plasmids and grown in 50 ml of LB media containing kanamycin (50 μg/ml), streptomycin (50 μg/ml) and chloramphenicol (34.5 μg/ml) overnight at 37°C shaking at 200 rpms in a 250ml Erlenmeyer flask. Next day, the bacterial culture was diluted (1:100) in LB media with the same antibiotic concentrations and grown at 37 °C until the bacterial culture reached an optic density of 0.15 (λ 600nm), then the culture was cooled to 18°C and tetracycline was added at a final concentration of 1ng/ml in order to trigger chaperone expression. When the optical density reached 0.4-0.5 (λ 600 nm), IPTG was added to a final concentration of 500 μM and culture was left overnight shaking (200 rpms) at 18°C. To express recombinant dSrc42A we followed a similar in which a pET-His10-Src42A plasmid was co-transformed with the pCFDUET-YopH construct only in BL21. Next day, bacterial culture was harvested at 3000 rpms and 4°C. The pellet was resuspended in 50 ml lysis buffer (50 mM Tris pH 8, 500 mM NaCl, 0.1mM PMSF) and sonicated on ice. Crude lysate was clarified by centrifugation at 20.000 rpms at 4°C for 45 min. Soluble clarified supernatant underwent a further 10 seconds sonication step and was filtered through a 45 μm syringe filter. Human c-Src was purified by three chromatographic steps (see figure 1). First, lysate was passed through an immobilized metal affinity chromatography (IMAC) column (GE Healthcare HisTrap™ HP) equilibrated with 5 column volumes (CVs) of IMAC buffer A (20 mM Tris pH 8, 150 mM NaCl, 1 mM TCEP, 5% Glycerol) with an GE Healthcare AKTA PURE FPLC at flowrate of 5 ml/min. Next, the column was washed with 90% IMAC buffer A and 10% IMAC buffer B (20 mM Tris pH 8, 150 mM NaCl, 300 mM Imidazole, 1mM TCEP, 5% Glycerol) until absorbance signal at a λ 280 nm was stable (usually 10 CVs), after which a 100% gradient with IMAC buffer B was run in 100 ml (20 CVs). Fractions containing the recombinant protein were collected and tested by SDS-PAGE before diluting in IEC (ionic exchange chromatography) buffer A (20 mM Tris pH 8, 1 mM DTT, 5% glycerol) up to 3 times original volume so NaCl concentration was lowered to 50 mM. Then, the sample was loaded into an GE Healthcare HiTrap Q HP column previously equilibrated with 5 CVs of IEC buffer A, then the column was washed with 5 CVs IEC buffer A, followed by and a 100% gradient with IEC buffer B (20 mM Tris pH 8, 500 mM NaCl, 1 mM DTT, 5% glycerol). Fractions containing the recombinant protein were collected and tested by SDS-PAGE, pulled together and mixed with a His-tagged rTEV protease (20-40 μM) in a 1/20 molar TEV/c-Src stoichiometry. Buffer was supplemented with 2 mM TCEP and digested at 4°C o/n. Alternatively, digestion was performed for 2 hours at room temperature. Next, a His-trap reverse step was undertaken to remove the protease and tag of the recombinant protein. Briefly, His-rTEV protease digested c-Src sample (input) was passed through the HisTrap column previously equilibrated with IMAC buffer A at 2 ml/min flowrate with a peristaltic pump (GE Healthcare Pump P-1) and then washed with 5 CVs of buffer IMAC A supplemented with 50 mM imidazole. Flow through (FT) was taken, checked by SDS-PAGE and UV-absorbance gel and (Thermo Scientific NanoDrop One) respectively. FT was taken, checked by SDS-PAGE and concentrated with a 10-30 kDa cut-off concentrator (Millipore) at 3.500 rpms at 4 °C. The sample was then injected in a size exclusion chromatography (SEC) Superdex 200 16/60 column (GE Healthcare) previously equilibrated with 1.5 CVs of SEC buffer (20 mM Tris pH, 150 mM NaCl, 1 mM DTT, 5% Glycerol). Fractions were collected and protein purity and concentration were tested by SDS-PAGE gel and UV-absorbance.

**Figure 1.**
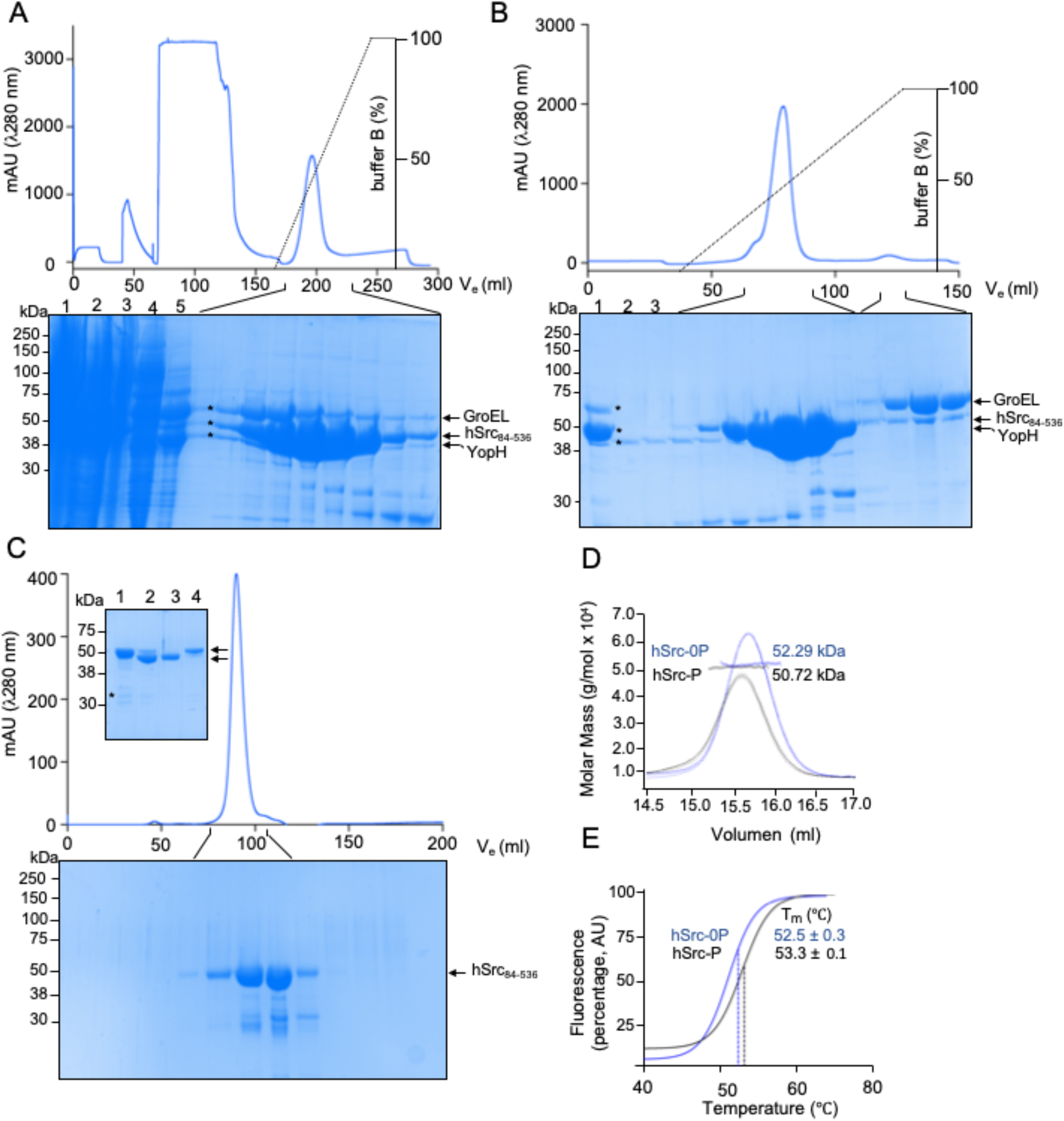
A tripartite expression system for heterologous production of c-Src in bacteria. (A) IMAC chromatogram using a HisTrap column (5 ml). Indicated fractions were run on an SDS-PAGE and stained with Coomassie: 1 crude lysate, 2 insoluble fraction, 3 clear lysate, 4 wash, 5 flow-through. (B) IEC chromatogram using a HiTrap Q HP column (5ml). Indicated fractions were run on an SDS-PAGE and stained with Coomassie: 1 pulled fractions from A, 2 wash and 3 flow-through. (C) Inset, HisTrap-reverse step. Indicated fractions were run on an SDS-PAGE and stained with Coomassie: (1) input from (B) i.e. pull of c-Src positive fractions from IEC, (2) pull of c-Src positive fractions after protease treatment, (3) flow-through and (4) elution with 500 mM imidazole. SEC chromatogram using a Superdex 200 16/60 column. Indicated fractions were run on an SDS-PAGE and stained with Coomassie. (D) SEC-MALS chromatogram showing UV (λ 280 nm) solid line, light scattering (dashed line), Rayleigh index (dashed soft line) and molar mass (g/mol x 10^4^) dots of un-phosphorylated c-Src (0-P, blue) and phosphorylated c-Src (P, 90 min, black). (E) DSF profile showing the melting temperature of un-phosphorylated c-Src (0-P) and phosphorylated c-Src (P, 90 min). Inset, data is the mean ± SEM of 4 experiments (n=4).

### Size-Exclusion Chromatography with Multi-Angle Light Scattering (SEC-MALS)

A protein sample (0.2 mg/ml) was injected in a Superdex 200 Increase 10/300 column (Cytiva) previously equilibrated in 20 mM Tris pH, 150 mM NaCl, 1 mM DTT, 5% Glycerol buffer (filtered through a 0.1 μm filter). The chromatographic eluent was monitored by three consecutive detectors in series: (1) a multi-wavelength UV-Vis absorbance detector Monitor UV-900 of the AKTA system (GE Healthcare) with a 10 mm path length flow cell, (2) a light scattering DAWN Heleos 8+ (Wyatt Technology) with detectors at eight different angles (from 32 to 141 degrees from the source) using a linearly polarized GaAs laser operating at 665 nm and (3) an Optilab T-rEX (Wyatt Technology) differential refractive index detector with a laser wavelength of 658 nm. Data collection and analysis were performed using UNICORN 5.10 (GE Healthcare) and ASTRA 6.0.3 (Wyatt Technology) software packages.

### Western blotting and antibodies

SDS-PAGE gels were transferred onto nitrocellulose 0.2 μm membranes (Amersham™). Transferred membranes were immersed in blocking solution (10 mM Tris pH 8, 150 mM NaCl, 5% weight/volume (w/v) skimmed powder milk) for 60 min. After blocking membranes were washed three times with TBS-T (10 mM Tris pH 8, 150 mM NaCl, Tween-20 0.1% v/v) prior incubation with primary antibody solution (TBS-T with BSA 5% w/v) o/n at 4°C shaking. Antibodies used were: phospho-Src Tyr419 (D49G4, CST #6943), Src (36D10, CST #2109) phospho-Src Tyr 530 (ThermoFisher 44-662G) and total phospho-Tyr (p-Tyr-100 CST #9411) were diluted at 1:10000-1:5000. After incubation with the first antibody, membranes were washed with 20 ml TBS-T three times and immersed in secondary antibody solution (TBS-T skimmed powder milk 5% m/v). Secondary antibodies anti-rabbit or -mouse IgG (DyLight conjugate at 680 or 800 nm (CST #5366, #5151, #5470, #5257) were used at half the dilution factor o the primary for one hour at room temperature protected from light. After incubation with secondary antibodies, membranes were washes 3-to-5 times with TBS-T. Membranes were scanned in an Odyssey CLx scanner.

### In vitro phosphorylation assays

In vitro phosphorylation assays were performed at room temperature using recombinant proteins at 1 μM final concentration in buffer (20 mM Tris pH, 150 mM NaCl, 1mM DTT, 5% Glycerol, 2 mM MgCl_2_) and ATP (1 mM) at the indicated time points (unstimulated, 1, 5, 15, 30, 60 and 90 min). Aliquots from the time-course were mixed with 5X Laemmli sample buffer (ThermoFisher) and denaturalized at 95°C during 1-2 min at 95 °C.

### Enzymatic assays

Phosphorylation rates of peptide substrates by recombinant proteins at 1μM final concentration were determined by using an NADH-coupled pyruvate kinase assay in presence of increasing ATP concentrations. The enzyme-substrate solution (20 mM Tris-Cl, 1 mM MgSO_4_, 400 mM Phosphoenolpyruvate (PEP), 100 mM NADH, 2450U/ml Pyruvate kinase (PK), 2.26 mg/ml Lactate dehydrogenase (LDH) was prepared at different concentrations of ATP: 0, 0.08, 0.16, 0.32, 0.65, 1.25, 2.5 and 5 mM. The experiments were performed in a 384 well plate (Greiner bio-one) and the NADH consumption was read at λ 340 nm during 120 cycles of 30 sec each by using a Victor multilabel plate reader 1420 Multilabel Counter (Perkin Elmer). In order to obtain catalytic rates and kinetic constants by Michaelis-Menten equation, experiments were analyzed using Prism software.

### Mass Spectrometry

For in-solution digestion, proteins samples (2-5 µM) were reduced and alkylated (15 mM TCEP, 30 mM CAA, 30min at RT in the dark) in the presence of urea 4 M and digested with trypsin in urea 1 M, 50 mM Tris pH 8 overnight at 37 °C (Promega) at an estimated protein:enzyme ratio 1:100. For FASP digestion, proteins were reduced and alkylated (15 mM TCEP, 30 mM CAA, 30 min in the dark, RT) and sequentially digested with chymotrypsin (SigmaAldrich) (protein:enzyme ratio 1:100, o/n at RT). In all cases, digestion was quenched by adding 0.1% TFA and resulting peptides were desalted using C18 stage-tips. LC-MS/MS was done by coupling an UltiMate 3000 RSLCnano LC system to a Q Exactive Plus mass spectrometer (Thermo Fisher Scientific). Peptides (2-5 µl) were loaded into a trap column (Acclaim™ PepMap™ 100 C18 LC Columns 5 µm, 20 mm length) for 3 min at a flow rate of 10 µl/min in 0.1% formic acid. Then, peptides were transferred to an EASY-Spray PepMap RSLC C18 column (Thermo) (2 µm, 75 µm x 50 cm) operated at 45 °C and separated using a 60 min effective gradient (buffer A: 0.1% FA; buffer B: 100% ACN, 0.1% FA) at a flow rate of 250 nL/min. The gradient used was, from 4% to 6% B in 2 min, from 6% to 33% B in 58 minutes, plus 10 additional minutes at 98% B. Peptides were sprayed at 1.5 kV into the mass spectrometer via the EASY-Spray source. The capillary temperature was set to 300 °C. The mass spectrometer was operated in a data-dependent mode, with an automatic switch between MS and MS/MS scans using a top 15 method (intensity threshold ≥ 4.5e4, dynamic exclusion of 5 or 10 sec and excluding charges unassigned, +1 and > +6). MS spectra were acquired from 350 to 1500 m/z with a resolution of 70,000 FWHM (200 m/z). Ion peptides were isolated using a 2.0 Th window and fragmented using higher-energy collisional dissociation (HCD) with a normalized collision energy of 27. MS/MS spectra resolution was set to 35,000 (200 m/z). The ion target values were 3e6 for MS (maximum IT of 25 ms) and 1e5 for MS/MS (maximum IT of 110 msec). Raw files were processed with Maxquant (v 1.6 and higher) using the standard settings against the corresponding sequences of the expressed recombinant proteins, and an E. coli protein database (UniProtKB/Swiss-Prot, 20,373 sequences) supplemented with contaminants. Carbamidomethylation of cysteines was set as a fixed modification whereas oxidation of methionines, protein N-term acetylation and phosphorylation of serines, threonines and tyrosines were set as variable modifications. Minimal peptide length was set to 7 amino acids and a maximum of two tryptic missed-cleavages were allowed. Results were filtered at 0.01 FDR (peptide and protein level). Raw data were imported into Skyline. Label free quantification of identified phosphopeptides was performed using the extracted ion chromatogram of the isotopic distribution. Only peaks without interference were used for quantification. Phosphopeptides intensities were normalized by the intensity of non-modified peptides from the target protein.

### Differential Scanning Fluorimetry (DSF)

To evaluate the thermal stability of recombinant c-Src in the absence of (apo) and in complex with Ponatinib, we applied an indirect SYPRO Orange-based method. For this assay the total reaction volume was adjusted to 40 µL at 1-2 µM protein, 10 µM inhibitor, and 2 x SYPRO Orange concentrations subjected to a gradient of temperature from 20 to 95 °C. Fluorescence was measured on an Applied Biosystem 7300 Real-Time PCR system.

### Drosophila work

Drosophila melanogaster strains were maintained and raised at 25°C under standard conditions. The stocks used are the following ones: GMRGal4 (eye expression, BSDC 9146), salEPvGal4 (wing expression, BDSC 80573), btlGal4 (tracheal expression, gift of S. Hayashi), src^F80^/CyOlacZ; UASSrc/TM6 dfdYFP (gift of S. Luschnig); UASSrcY400F/TM6 dfdYFP (this work). Balancer chromosomes CyO, TM3 or TM6 marked with LacZ, GFP or YFP were used to follow the mutations and constructs of interest in the different chromosomes. Transgenes were generated by injecting the pUAST-attB-Src42a WT and Y400F construct to obtain directed insertions at 68E by the “Transgenesis Service” of the “Centro de Biología Molecular Severo Ochoa” (CBM, Madrid). Transgene expression was achieved using the Gal4/UAS system (Brand and Perrimon, 1993) at 25°C (GMRGal4) or 29°C (btlGal4 and salEPvGal4). Embryos were stained following standard protocols. Primary antibodies used were goat anti-GFP (1:600) from Roche and chicken anti-β-gal (1:200) from Abcam. Alexa Fluor 488, 555, 647 (Invitrogen) secondary antibodies were used at 1:300 in PBT 0.5% BSA. CBP (Chitin Binding Protein, produced by N. Martín in Dr. Casanova’s lab, New England Biolabs Protocol) was used as a secondary antibody at 1:300 to detect chitin and visualize the tracheal branches. Images from fixed embryos were taken using Leica TCS-SPE with the 20x and 63x immersion oil (1.40-0.60; Immersol 518F – Zeiss oil) objectives and additional zoom. Images from adults were obtained with an Olympus MVX10 macroscope using EFI (extended focus imaging) at the ADM facility of IRB-PCB. For morphometric analyses confocal projections were used to analyze the length of the embryonic dorsal trunk (DT) stained with CBP in stage 16 embryos. We traced the path using the freehand line selection tool of Fiji (ImageJ) software between the junction DT/Transverse Connective (TC) from metamere 2 to 9 following the DT curvature. DT length was expressed as the ratio between the DT path and the length of the embryo. Data from quantifications was imported and treated in the Excel software and in GraphPad Prism 9.0.0, where graphics were finally generated. Graphics shown are scatter dot plots, where bars indicate the mean and the standard deviation. Statistical analyses comparing the mean of two groups of quantitative continuous data were performed in GraphPad Prism 9.0.0 using unpaired two-tailed student’s t-test applying Welch’s correction. Differences were considered significant when p < 0.05. *p < 0.05, **p < 0.01, *** p < 0.001, **** p < 0.0001 where n.s. means not statistically significant.

### Crystallization, Diffraction, Data Collection, and Processing

Crystals of the c-Src KD (aa 252-536, human) in complex with ponatinib were obtained by mixing recombinant protein and ponatinib in a 1:2 molar ratio. Sample was concentrated using an Amicon Ultra 10K molecular weight cut-off centrifugal filter device (Merck Millipore, Billerica, Massachusetts, USA) up to 8 mg/ml. The final protein concentration was determined by UV spectroscopy (Nanodrop one, Thermo Scientific). Crystallization for c-Src KD in complex with ponatinib was achieved by sitting-drop vapour diffusion at 20 °C in MRC-2 crystallization plates (Molecular Dimensions, Newmarket, Suffolk, England) by mixing 0.5 μL of protein-ligand solution at 8 mg/ml with 0.5 μL of reservoir solution. After several days some crystals were obtained in a drop containing 50 mM sodium acetate pH 4.6, 100 mm sodium chloride, 20% w/v PEG 4000 and 10% v/v 2-Propanol. A cryoprotectant solution consisting of the reservoir solution including 20% (v/v) glycerol was used to freeze the crystals and mounted in LithoLoops (Molecular Dimensions, Newmarket, England) prior to vitrification in liquid nitrogen for data collection. X-ray diffraction data were collected at a wavelength of 0.98 Ǻ on beamline XALOC-BL13 of the ALBA Synchrotron Light Facility (Barcelona, Spain) using a Pilatus 6M pixel detector (Dectris Ltd, Baden, Switzerland). Crystals were kept at 100 K during data collection. Reflections were integrated with the programme iMOSFLM and reduced using POINTLESS, AIMLESS and TRUNCATE, all integrated in the Collaborative Computational Project Number 4 (CCP4). Molecular replacement was carried out using the atomic coordinates of c-Src KD in complex with imatinib (PDB 3EL8) using PHASER (McCoy, 2007). Adjustment of the model was performed with COOT (Emsley et al., 2010) and refinement was carried out with Refmac5 (Murshudov et al., 2011) applying twin refinement amplitude-based settings. Model validation was carried out with MOLPROBITY and structure figures were made using the graphics program PYMOL. For data statistics, see supplemental table 1.

### Small angle X-ray scattering (SAXs)

SAXS experiments were conducted at the beamline B21 of the Diamond Light Source (Didcot, UK) (Cowieson et al., 2020). A sample of 40 ul of Src WT (3D-construct) at concentration of 3 mg/ml was delivered at 20°C via an in-line Agilent 1200 HPLC system in a Superdex 200 Increase 3.2 column, using a running buffer composed by 20 mM Tris pH 8.0, 150 mM NaCl, 1% glycerol and 1 mM DTT. The continuously eluting samples were exposed for 300s in 10s acquisition blocks using an X-ray wavelength of 1 Å, and a sample to detector (Eiger 4M) distance of 3.7 m. The data covered a momentum transfer range of 0.0032 < q < 0.34 Å-1. The frames recorded immediately before elution of the sample were subtracted from the protein scattering profiles. The Scåtter software package (www.bioisis.net) was used to analyse data, buffer-subtraction, scaling, merging and checking possible radiation damage of the samples. The R_g_ value was calculated with the Guinier approximation assuming that at very small angles q < 1.3/R_g_. The particle distance distribution, D_max_, was calculated from the scattering pattern with GNOM, and shape estimation was carried out with DAMMIF/DAMMIN, all these programs included in the ATSAS package (Petoukhov et al., 2012). The proteins molecular mass was estimated with GNOM. Interactively generated PDB-based homology models were made using the program COOT by manually adjusting the X-ray structures obtained in this work, into the envelope given by SAXS until a good correlation between the real-space scattering profile calculated for the homology model matched the experimental scattering data. This was computed with the program FoXS (Schneidman-Duhovny et al., 2016).

## RESULTS

### 1. A tripartite expression system for heterologous production of c-Src in bacteria

Biochemical characterization of recombinant protein kinases usually requires low amounts (nano-to-micrograms) of material. On the contrary, biophysical and X-ray crystallography studies demand milligram amounts of homogenous protein of high stability and purity. We have applied a tripartite expression system for the heterologous expression of c-Src in bacteria. The tripartite system allows the quick production and purification of high-yields of active and monodisperse c-Src by co-expression with the YopH tyrosine phosphatase and the GroEL/GroES chaperones. Our procedure is an updated version of a previously stablished protocol (Seeliger et al., 2005) where high-yield production of active and soluble c-Src KD required the co-expression of YopH phosphatase. One of the main caveats of this preliminary bi-partite system was that 90% of the protein was still found in the insoluble fraction. Addition of the GroEL chaperone into the system increased solubility by 5-10-fold (Fig. 1A). First, we expressed a three domain (3D, SH3-SH2-KD) construct (aa 84-536) in BL21 cells (see material and methods) and performed tandem immobilized metal affinity (Fig. 1A), cation exchange (Fig. 1B) and size exclusion (Fig. 1C) chromatography, in order to separate each individual component to obtain pure and monodisperse c-Src. We applied multi-angle light scattering (MALS) for determination of absolute molar mass in solution and found that both, the un-phosphorylated and phosphorylated forms of c-Src are active monomers in solution (Fig. 1D and Fig. S1A). In line with this, phosphorylation does not cause any significant effect on the thermal stability of the protein as indicated by differential scanning fluorometry (DSF) either (Fig. 1E). These data however do not exclude the fact that the intrinsically disordered N-terminal region lacking in the 3D construct could be implicated in the dimerization process (Spassov et al., 2018). To test this hypothesis, we expressed and purified a full-length construct of the drosophila c-Src homolog Src42A (see material and methods). We found again that the full-length protein is active (work in preparation) and monomeric (60 kDa) in solution, and that auto-phosphorylation does not induce direct dimerization (Fig. S1B and C). In light of these results, we explored further the potential role of myristoylation on c-Src function and stability by using native and G2-myristoylated peptides in activity and binding (DSF) assays (Fig. S1D). Enzymatic assays performed with c-Src 3D and KD constructs demonstrated that preincubation of the enzyme with peptide (1:2.5 molar ratio, 45 min) does not result in significant changes in the catalytic rates of both constructs toward an exogenous susbtrate. In DSF experiments (Fig. S1D) none of the peptides caused a significant increment in thermal stability that would have been otherwise indicative of efficient binding. These data together demonstrate that c-Src is an active monomer in solution and that auto-phosphorylation does not promote direct dimerization. Myristoylation on the intrinsically disordered N-terminal region appears not to affect the dimerization nor direct interaction with the catalytic domain in vitro, and lacking any regulatory effects on the catalytic activity.

### 2. Kinetics of c-Src auto-phosphorylation

In the current paradigm, c-Src is held in an autoinhibited state by intramolecular interactions between the SH2 domain and CSK-phosphorylated Tyr 530 on the c-terminal segment (Fig. 2A). To define the kinetics of c-Src intrinsic auto-phosphorylation in the absence of CSK we used a c-Src 3D-construct and performed in vitro time course auto-phosphorylation assays (0–90 min). Auto-phosphorylation was monitored by Western blotting (WB) using total and phospho-specific antibodies against Tyr 216, Tyr 419 and Tyr 530 (Fig. 2B). Activation loop Tyr 419 displays fast auto-phosphorylation kinetics reaching saturating levels at early time points (1 to 5 min). In contrast, c-terminal Tyr 530 and SH2 Tyr 216 show slower rates of auto-phosphorylation reaching maximum levels at later time points (30-60 min). Enzymatic assays using peptides derived from activation loop and c-terminal phospho-sites (Fig. 2C) corroborated the WB data, with a five-fold increase in the catalytic efficiency constant (k_cat_/K_M_) by the Tyr419-derived peptide over Tyr 530. Interestingly, these data highlight two important observations: i) the existence of sequential and coordinated auto-phosphorylation events coupled to different conformation states, and ii) c-terminal Tyr 530 is a de facto c-Src auto-phosphorylation site. Considering these two premises, the activation loop of c-Src must be readily accessible to be phosphorylated, while slower c-terminal Tyr 530 auto-phosphorylation occur at later stages due to restricted accessibility by coordination between the SH2 domain and (un-phosphorylated) Tyr 530 as shown by SAXS analysis (Fig. S2). These data corroborated that in solution un-phosphorylated c-Src 3D is adopting a closed state resembling the PDB 2SRC. Next, we used high-resolution mass spectrometry (MS) to monitor simultaneously multiple auto-phosphorylation sites. We identified twenty-five phospho-sites, among them some unexpected serine and threonine residues as well as new tyrosine sites (Fig. S3). Novel phospho-sites identified and validated in at least in two independent experiments were Tyr 95 (βa/βb linker, SH3), Tyr 152 (βA, SH2), Thr 288 (β2), Tyr 385 (αE/β7 linker) and T443 (αF) in the kinase domain (Xu et al., 1997). MS data on those best characterized phospho-sites validate our WB data using phospho-specific antibodies (Fig. 2C) where again quick kinetics of auto-phosphorylation for activation loop Tyr 419 contrasted with the slower rates of c-terminal Tyr 530 phosphorylation.

**Figure 2.**
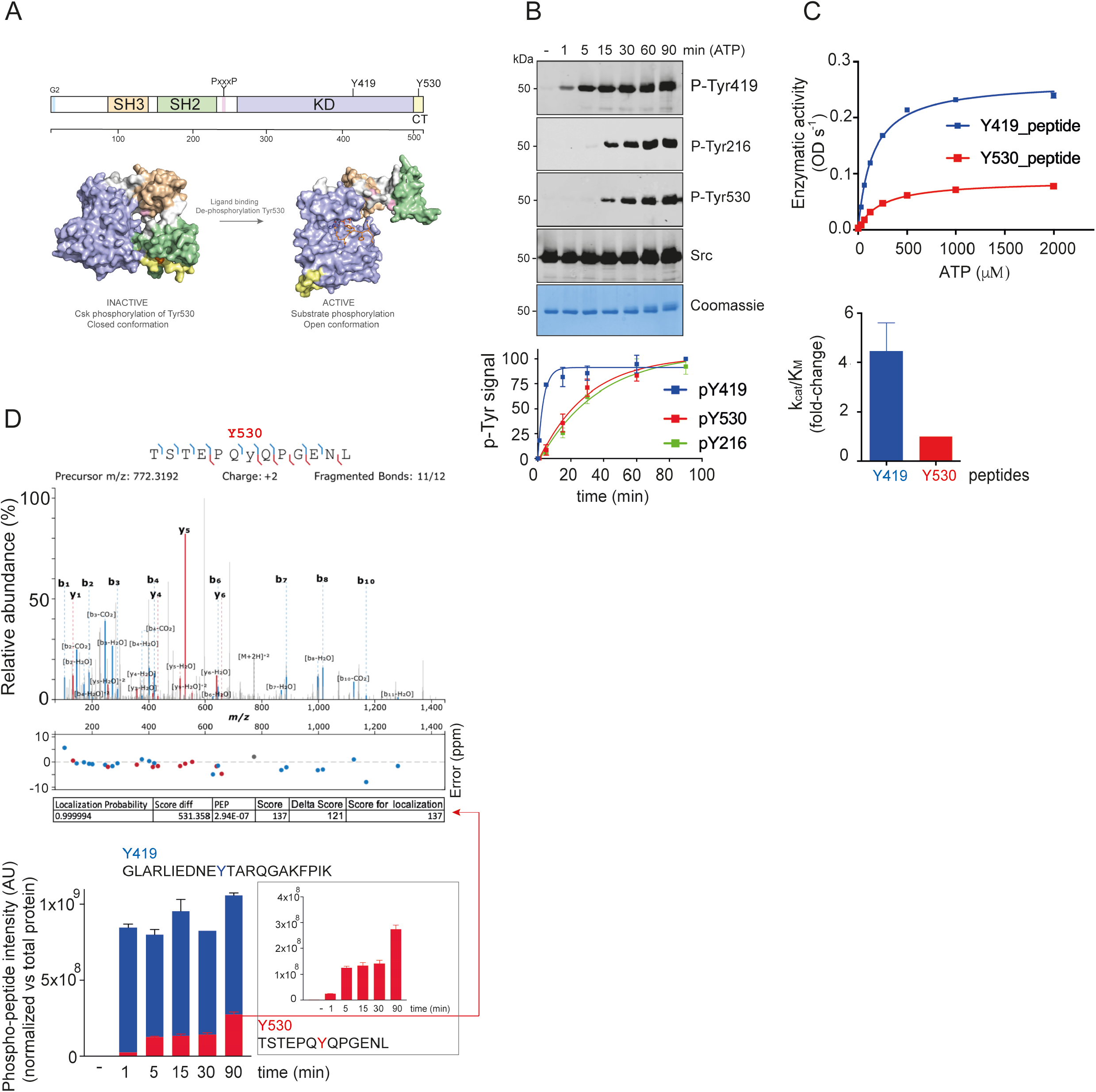
Kinetics of c-Src auto-phosphorylation. (A) Schematic diagram of the functional domains and auto-phosphorylation sites of c-Src. Surface representation of c-Src in closed autoinhibited and open (active) states, current paradigm for c-Src activation and regulation. (B) WB of samples from a time-course auto-phosphorylation experiment with c-Src WT (3D-construct, 1 μM) in the presence of ATP (1 mM) and MgCl_2_ (2 mM) for 0–90 min using the indicated antibodies. Total amount of protein was visualized by Coomassie staining. Phospho-tyrosine quantification (total), data represent mean ± SEM of 6 experiments (n=6). (C) Enzymatic assay performed with c-Src WT (3D-construct, 1 μM) incubated with increasing concentrations of ATP at a fixed concentration (1.5 mg/ml) of c-Src Y419 (IEDNEYTARQG) or Y530 (STEPQYQPGEN) derived peptides. Data represent the mean ± SEM, of 2 experiments (n=2) in duplicate. Enzymatic activity (ODs^-1^ x 10^−3^). Catalytic efficiency constants (k_cat_/K_M_, fold difference) are depicted in the panel below. (D) Mass spectrum of the [M + 2H] +2 ion (m/z 772.3) of a peptide phosphorylated on Tyr 530 (90 min). Below, phosphorylation kinetics of Tyr419 and Tyr530 phospho-peptides measured by mass spectrometry (0-90 min) are depicted. Data represent the mean of normalized phospho-signal ± SEM of 2 experiments (n=2), with two technical replicates each.

### 3. c-Src activation-loop Tyr 419 controls enzyme specificity

To evaluate *in vitro* the activating and inhibitory roles previously described in a cellular context for c-Src activation loop Tyr 419 and c-terminal Tyr 530 respectively (Roskoski, 2004, 2005), we generated tyrosine to phenylalanine mutants in a c-Src 3D-construct and performed in vitro time course auto-phosphorylation assays (Fig. 3A). In these experiments we observed a significant degree of phospho-tyrosine activity by a c-Src activation loop mutant Y419F, which was consistently lower compared with the WT. Remarkably, this mutant lacks the capacity to auto-phosphorylate on Tyr 530 at the c-tail (Fig. 3A). Conversely, a c-Src Y530F mutant displays higher phospho-tyrosine activity, which is also reflected in faster kinetics of Tyr 419 and Tyr 216 auto-phosphorylation compared with the WT c-Src (Fig. 3A). These results were further confirmed by MS (Fig. S4). Next, we performed enzymatic assays using a generic peptide derived from c-Abl as an exogenous substrate (Fig. 3B). Under these conditions, the three constructs efficiently phosphorylate the c-Abl peptide, with only a two-fold difference in the V_max_ observed between the WT and Y419F. On the contrary, when a peptide derived from the RET activation loop was tested, there was one order of magnitude difference in the V_max_ between Y419F and the WT (Fig. 3B). In these experiments, WT and c-terminal Y530F mutant displayed similar activities towards the exogenous substrate peptides. These data suggest changes in the susbtrate preferences (rather than catalytic impairment), by a non-phosphorylable c-Src activation loop mutant. To prove this hypothesis, we used a series of recombinant proteins as intact substrates in in vitro phosphorylation assays: a fragment of human FAK (aa 1-405) that contains Tyr 397, c-Src KD K298M, RET KD K758M and deltaKIF5B-RET construct (Fig. 3 C-F). All these proteins lack catalytically activity and hence act as substrate surrogates. c-Src 3D constructs WT, Y419F and Y530F were able to efficiently phosphorylate GST-FAK (aa1-405) and c-Src KD with no significant differences among them (Fig. 3C-D). Strikingly, when a RET KD was used as an intact susbtrate, c-Src Y419F contrary to WT or Y530 was not able to phosphorylate the activation loop on Tyr 905 as indicated by a phospho-specific RET antibody (Fig. 3E). These data were further corroborated using a truncated KIF5B-RET fusion that lacks also catalytic activity, acting as an intact susbtrate surrogate (Fig. 3F). In both cases, a c-Src Y419F mutant was unable to efficiently phosphorylate RET. Consensus phospho sites sequences scrutiny reveal that out of the three substrates surrogates used, the RET activation loop sequence was the most similar to the optimal c-Src substrate sequence (phosphosite.org), hence from these data we conclude that c-Src activation-loop Tyr 419 controls enzyme specificity towards substrates (Fig. 3G).

**Figure 3.**
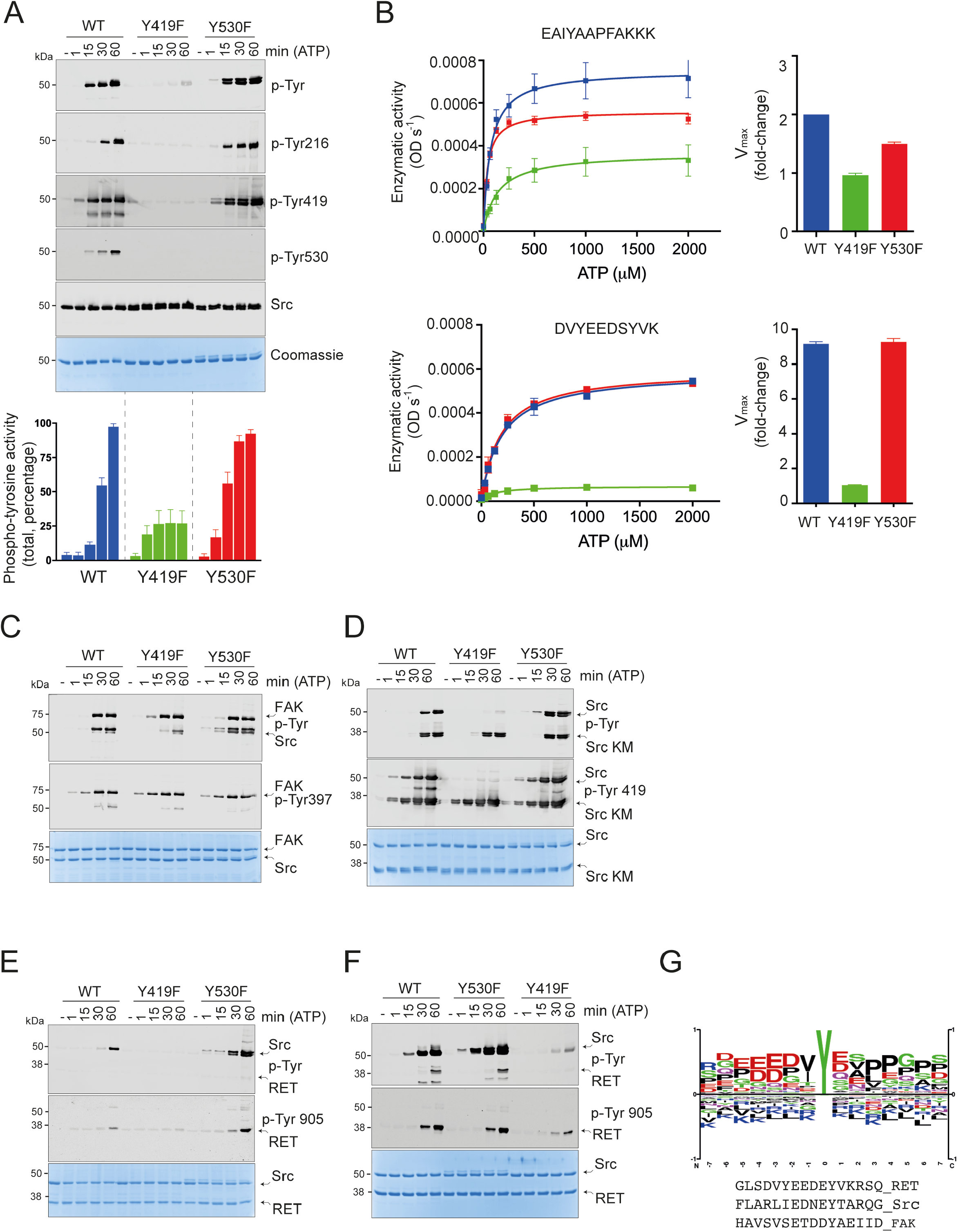
Activation-loop Tyr 419 controls c-Src susbtrate specificity. (A) WB of samples from a time-course auto-phosphorylation experiment with c-Src WT, Y419F and Y530F (3D-construct, 1 μM) in the presence of ATP (1 mM) and MgCl_2_ (2 mM) for 0–60 min using the indicated antibodies. Total amount of protein was visualized by Coomassie staining. Phospho-signal quantification (total phospho-tyrosine), data represent mean ± SEM of 6 experiments (n=6) in duplicate. (B) Enzymatic assay performed with c-Src WT, Y419F and Y530F (3D-construct, 1 μM) incubated with increasing concentrations of ATP at a fixed concentration (4 mg/ml) of peptide: ABL derived-peptide (EAIYAAPFAKKK) or RET activation loop Y905 derived-peptide (DVYEEDSFVK). Data represent the mean ± SEM, of 4-6 experiments (n =4-6) in duplicate. Catalytic efficiency constants (k_cat_/K_M_, fold difference) are depicted in the right panel. (C-F) WB of samples from a time course phosphorylation assays with c-Src (3D, 1 μM) WT, Y419F and Y530F and using the following substrates surrogates: FAK (aa 1-405), c-Src KD K298M, RET KD (aa 713-1012) K758M and deltaKIF5B-RET, respectively using the indicated antibodies. (G) Logo consensus sequence for optimal c-Src susbtrate, from PhosphoSitePlus (CST).

### 4. Time resolved intra- and -inter-molecular components for the mechanism of c-Src auto-phosphorylation

In order to dissect the time-resolution of the inter- and intra-molecular components for the mechanisms of c-Src Tyr 419 auto-phosphorylation, we performed time course reactions with increasing protein concentration (0.25, 0.5, 1 and 2 μM) of c-Src 3D WT, Y419F and Y530F. Equal amounts of samples (see Coomassie) were loaded and WB performed using phospho-specific antibodies (Fig. 4A). In the case of c-Src WT, we observed a fast cis-component (protein concentration independent) for Tyr 419 auto-phosphorylation at early time points, e.g. at 1 min see 0.25, 0.5 and 1 μM concentrations and at 5 min, see 0.25 and 0.5 μM concentrations. This was followed by an intermolecular component (protein concentration dependent) at later time stages e.g. see 15 min at 0.5 μM, 5 min at 1 μM and 1 min at 2μM concentrations (Fig. 4A). On the other hand, c-terminal Tyr 530 auto-phosphorylation showed a strong intermolecular component (see figure 6D). Strikingly, activation loop Tyr 419 phosphorylation by a c-terminal Y530F mutant showed a very strong cis-component as indicated by very fast kinetics of Tyr 419 auto-phosphorylation, which were independent of enzyme concentration (8-fold increase, from 0.25 to 2 μM). Altogether these data showed that in the native state the mechanism of c-Src activation loop auto-phosphorylation is defined by a fast intra-molecular component that switches on time to an inter-molecular process, i.e. a cis-to-trans auto-phosphorylation switch. Furthermore, the crosstalk between the activation-loop and c-terminal segment is coordinated and coupled to different conformational and functional states as indicated by: i) the lack of inter-molecular component by a mutant that cannot be phosphorylated on Tyr 530, and ii) an activation loop mutant cannot phosphorylate c-terminal Tyr 530. Next, we evaluated the effect on enzyme concentration (from 0.25 up to 2 μM) in the kinetics rates and constants of c-Src WT, Y419F and Y530F for ATP using a c-Abl derived peptide (Fig. S5). Src WT showed constant V_max_ and k_cat_ but lower K_M_ values, with an 8-9-fold increase in the catalytic efficiency constant over the range of enzyme concentrations used. Whereas c-Src Y419F showed increased V_max_ and constant K_M_ for ATP, c-Src Y530F mutant showed a constant V_max_ and a significant decreased in K_M_ that resulted in a significant 20-fold increase in the catalytic efficiency constant (k_cat_/K_M_) over the range of enzyme concentrations. These data altogether demonstrate that c-Src activation loop mutant Y419F is an active enzyme, and that Src c-terminal Tyr 530 mutant displayed increased catalytic efficiency constant for ATP compared with WT and Y419F, which support the results from the autophosphorylation assays, where c-Src Y530F displayed higher phospho-tyrosine activity. Next, in order to discriminate the catalytic (enzyme) from non-catalytic (susbtrate) properties we used first a c-Src KD K298M catalytically impaired mutant as an intact substrate surrogate in phosphorylation assays (Fig. 4B). In this system, c-Src WT and Y530F efficiently phosphorylated the activation loop of the susbtrate. Unexpectedly, a c-Src Y419F mutant did also efficiently phosphorylate the susbtrate that mimics its own activation loop, however and contrary to the WT and Y530F mutant, it did not phosphorylate c-terminal Tyr530 in trans. These results further confirm that activation loop Tyr 419 is required for c-terminal Tyr 530 phosphorylation, and that none of the two main auto-phosphorylation sites are required to maintain the enzymatic activity of c-Src.

**Figure 4.**
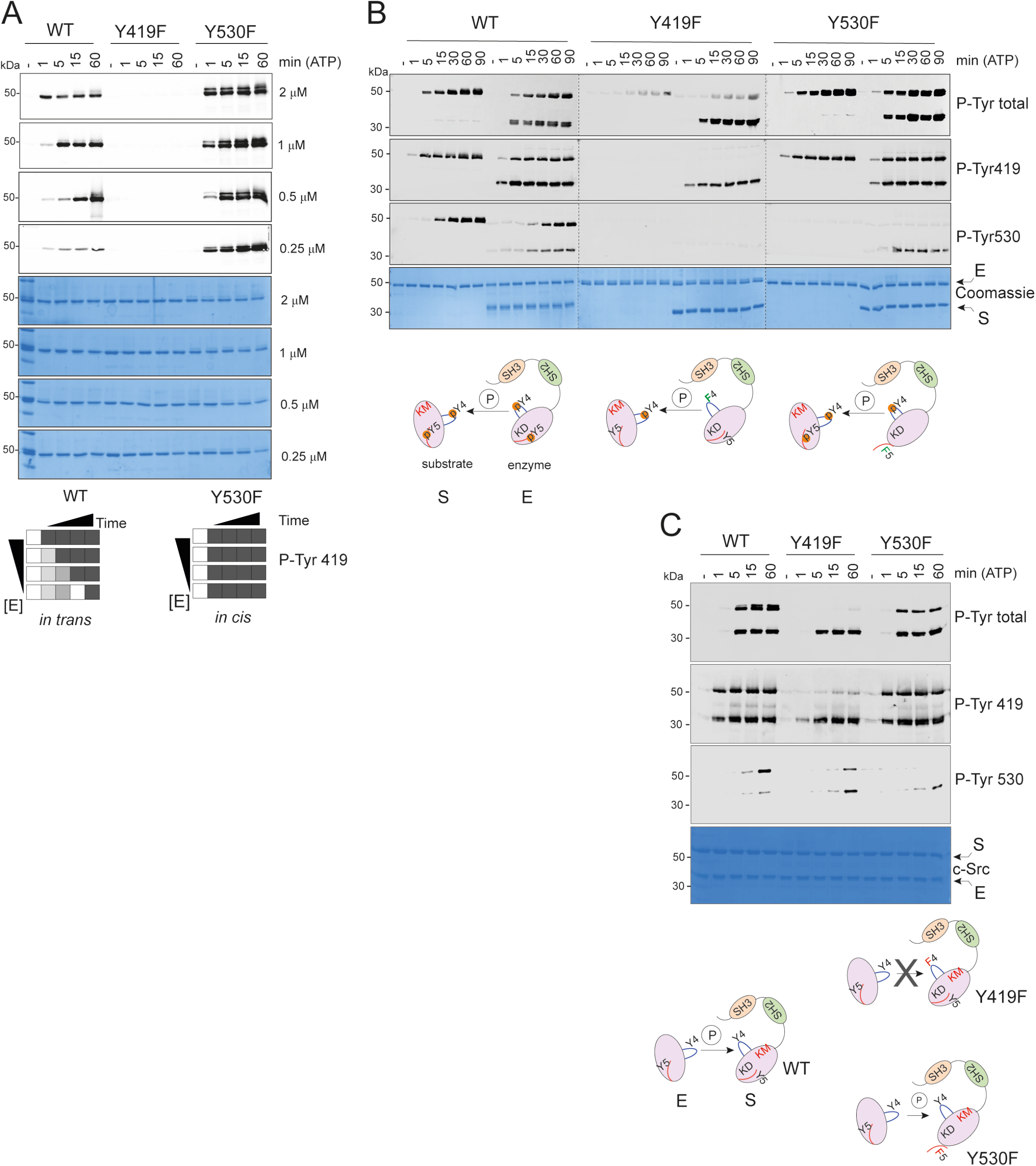
Dissecting cis-versus-trans components for c-Src auto-phosphorylation. (A) WB of samples from a time-course auto-phosphorylation experiment with WT, Y419F and Y530F c-Src (3D-construct, 0.25-2.5 μM) in the presence of ATP (1 mM) and MgCl_2_ (2 mM) for 0–60 min using the indicated c-Src phospho-Tyr 419 antibody. Total amount of protein was visualized by Coomassie staining. Diagram for the in-trans and cis-mechanism for auto-phosphorylation, lower inset. (B) WB of samples from a time-course phosphorylation assay (0-90 min) as in A, in the absence and in the presence of a c-Src KD K298M construct as an intact susbtrate surrogate using the indicated antibodies. Total amount of protein was visualized by Coomassie staining. Diagram for the enzyme (E) and susbtrate (S) acting kinases. lower inset. (C) WB of samples from a time-course experiment with c-Src KD WT in the presence of c-Src 3D-K298M constructs with and without Y419F and Y530F mutations as substrates using the indicated antibodies. Total c-Src protein was visualized by Coomassie staining. Diagram for the enzyme (E) and susbtrate (S) acting kinases. lower inset.

### 5. Activation-loop Tyr 419 controls c-Src non-catalytic properties

Second, to explore the non-catalytic (susbtrate) functions of c-Src we performed the opposite time-course phosphorylation experiments using a c-Src KD construct as an active enzyme and a catalytically impaired 3D-K298M construct with and without activation-loop Y419F or c-terminal Y530F mutations as substrate surrogates (Fig. 4C). In the case of a K298M construct, while activation-loop Tyr 419 is efficiently phosphorylated by the enzyme acting kinase, c-terminal Tyr 530 appears to be a less efficient phosphorylable substrate. These data come in line with the preferences observed in the dynamics of auto-phosphorylation and enzyme kinetics data using phospho-specific antibodies and peptides derived from each consensus phospho-sites (Fig. 2 A and C). Strikingly, a susbtrate surrogate with a Y419F mutation at the activation loop behaves overall as a very poor substrate, showing neglected phosphorylation promoted by the enzyme acting kinase (Fig 4C). A substrate with a non-phosphorylable c-terminal Tyr 530 is efficiently phosphorylated by an active kinase molecule, however up to a lower level (2-fold) compared with the native substrate (Fig 4C). These results demonstrate that Tyr 419 controls in addition to substrate specificity, important susbtrate-like non-catalytic properties by means of presenting the activation loop to another active moleculefor intermolecular phosphorylation.

### 6. A *Drosophila* model for the in vivo evaluation of a c-Src mutant at the activation loop

To understand the physiological activity of activation loop Tyr 419 in an in vivo system we used the fruit fly *Drosophila* melanogaster. We generated transgenic lines expressing the Src42A Y400F mutation (equivalent to Tyr 419 in human c-Src) in different tissues (Fig. 5). First, we compared the phenotypic effects produced by the expression of Src42A Y400F with the phenotype produced by the expression of a WT allele (Fig. 5A). When Src42A WT was expressed in the eye by means of the Gal4/UAS system using the GMR promoter, we observed a strong phenotype in the eye in which flies virtually lack all ommatidia (Fig. 5A), as previously described (Pedraza et al., 2004; Putz, 2019). The expression of Src42A Y400F in the eye also produced a clear and penetrant rough eye phenotype, however the defects were weaker than those observed with the WT protein (Fig. 5A). This result indicated that Src42A Y400F is functional, although likely less active than the Src42A WT. In the same line, expression of Src42A Y400F in the wing using a SalEPV-Gal4 driver resulted in a penetrant phenotype of rudimentary wings (Fig. 5B). Again, expression of Src42A Y400F produces a dominant effect but yet weaker than the one observed when expressing WT, indicating that the protein is active but may lack some particular functional facet.

**Figure 5.**
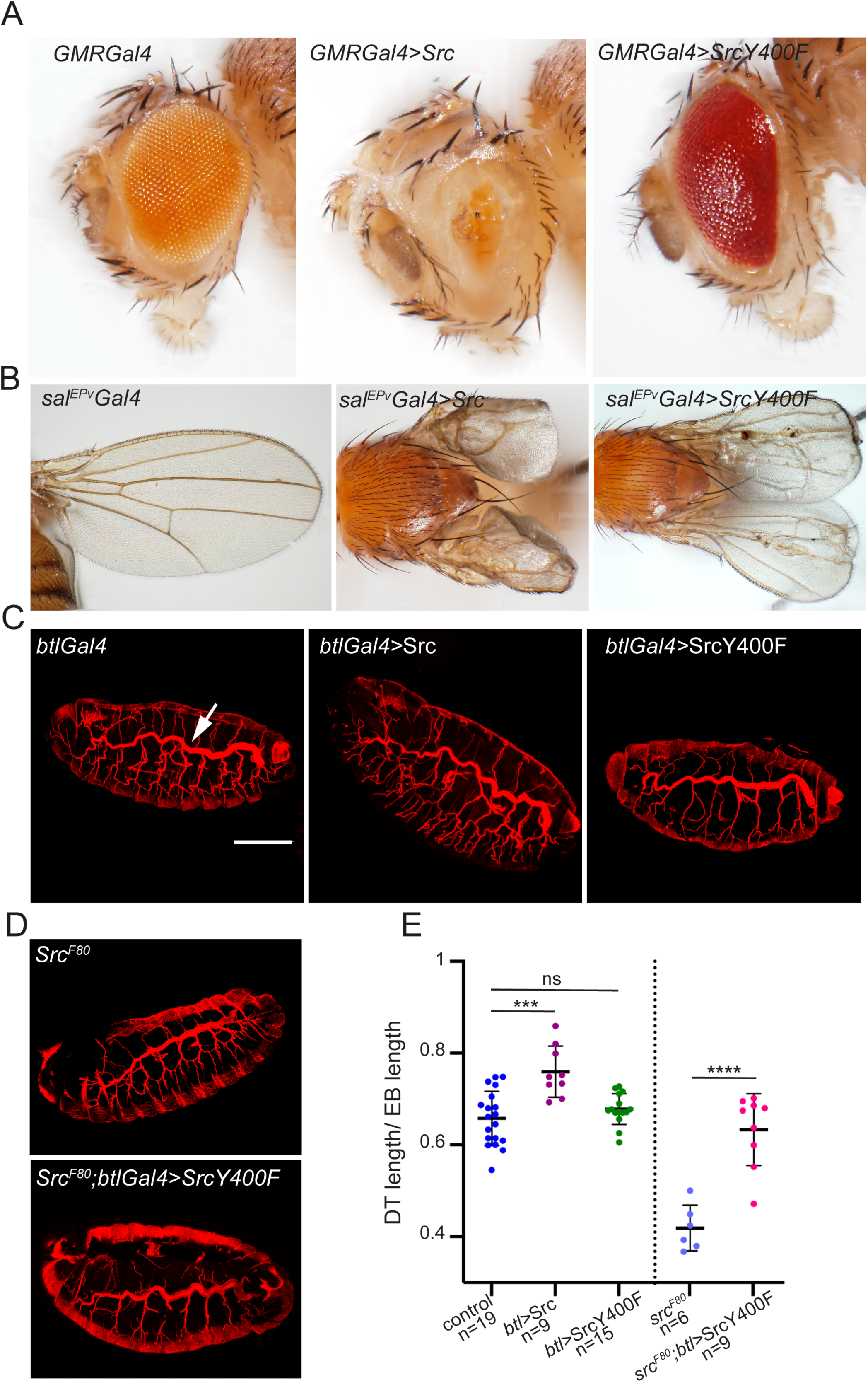
In vivo evaluation of a c-Src mutant at the activation loop using *Drosophila* (A) Expression Src42A WT and Y400F mutant in the *Drosophila* eye using a GMR-Gal4 promoter. Representative images of eyes from GMR-Gal4 (control), GMR-Gal4-Src42A WT and GMR-Gal4-Src42A Y400F flies. (B) Expression of Src42A WT and Y400F in the *Drosophila* wing using a salEPv-Gal4 promoter. Representative images of wings from salEPv-Gal4 (control), salEPv-Gal4-Src42A and salEPv-Gal4-Src42A Y400F flies. (C) Expression of Src42A WT and Src42A Y400F in the developing *Drosophila* tracheal tube using the btl-Gal4 promoter. Images show projections of confocal sections of lateral views of stage 16 embryos stained for CBP to visualize the tracheal tubes. The length of the DT (arrow in control) is quantified in E. Scale bar, 100 μm (D) Rescue experiments to assay Src42A Y400F function. Projections of confocal sections of lateral views of stage 16 embryos stained for CBP to visualize the tracheal tubes. *Src* mutant embryos (*Src*^*F80*^) show reduced DT length. Expression of Src42A Y400F in the trachea of *Src*^*F80*^ mutants restores DT length (quantifications are shown in E).

Src42A has been shown to play a key role during the embryonic development of the tracheal (respiratory) system. In particular, Src42A is required for the adequate orientation of the membrane growth along the longitudinal axis of the tracheal tubes regulating in this way their size (Forster and Luschnig, 2012; Nelson et al., 2012). Next, to investigate the phenotypic impact of Src42A Y400F in the developmental trachea we applied the Gal4/UAS system with the breathless (btl) promoter (Fig. 5C) in an intact genetic background. While the lack of Src42A (SrcF80) activity lead to shorter tubes, expression of Src42A WT leads to a moderate elongation of the tracheal tubes in larvae (Forster and Luschnig, 2012). In contrast, we found that the expression of Src42A Y400F resulted in a lack of significant elongation (or reduction) of the tube size (Fig. 5C). These data suggest that an activation loop mutant Src42A protein, although functionally competent, is less efficient and contrary to the expression of a native protein results in a lack of dominant effect in the trachea. Alternatively, it may indicate that the mutated protein is not functional in the trachea. To discriminate between these two possibilities, we expressed Src42A Y400F in a loss of function genetic background for Src using SrcF80 flies (Nelson et al., 2012). In this experimental setting, we observed that Src42A Y400F is able to fully rescue the tracheal defects seen in Src F80 flies, reverting the defects of short tubes, as previously shown for Src42A WT (Nelson et al., 2012). These results demonstrate that Src42A Y400F is functional and capable of rescuing the developmental defects promoted by a loss of function Src42A allele in the trachea.

### 7. Crystal structure of c-Src KD in complex with Ponatinib

We solved the crystal structure of a c-Src KD construct (aa 252-536) in complex with Ponatinib at 2.49 Å resolution (Fig. S6A). The crystal belonged to the orthorhombic system, and data was processed using P1211 symmetry (Table S1). The crystal had two molecules of c-Src in the asymmetric unit, and in both chains the electron density for Ponatinib was perfectly defined with full-occupancy. The central part of the activation loop (aa 410-425) was disordered and lacked electron density (Fig. S6A and B). Ponatinib binds to the active site in an extended pose reaching the back and allosteric pockets making the activation loop to adopt a DFG-out conformation (Fig. S6A-C). This configuration was also seen in the crystal structures of c-Src, c-Abl and c-Kit in complex with Imatinib (Mol et al., 2004; Nagar et al., 2002; Schindler et al., 2000; Seeliger et al., 2007). The imidazopyridazine core of Ponatinib rested on the adenine pocket of the enzyme and formed hydrophobic interactions with residues located within and at the vicinity of the hinge A296 (β3), L276 (β1), L396 (β6) and Y343 (hinge), forming one hydrogen bond with main chain nitrogen of M344 (hinge). The ethynyl linkage forms favorable hydrophobic interactions and a halogen bond with V326 (αC-β4 loop). The methylphenyl group occupies a hydrophobic pocket formed by T341 (β4) and I339 (β4), with the oxygen atom from the methyl group forming an additional hydrogen bond with the catalytic K298 (β3). The trifluoromethyl group forms hydrophobic interactions with A406 (A-loop), V405 (A-loop) and also to L325 (αC-β4 loop), while the phenyl group forms hydrophobic contacts with M317 (αC). The methyl piperazine group formed electrostatic interaction with D407 (A-loop) inducing altogether a DFG-out conformation (Fig. S6A-D) presenting an additional hydrogen bond with main chain oxygen of H387. The DFG-out conformation is not compatible with the proper alignment of the regulatory R-spine (Kornev et al., 2006), which is broken by the flipping of the DFG-motif, which is a signature of inactive activation loop conformation (Fig. S6E).

### 8. A crystallographic snapshot of c-terminal Tyr 530 intermolecular phosphorylation

Another interesting feature of the crystal structure presented in this study is the arrangement of the two molecules of c-Src found in the asymmetric unit, where the back side of the c-lobe of one protomer faced the susbtrate binding area located at the base of the active site cleft of the other (Fig. 6A and B). The interface is composed by the αI-I′-helix (residues 512-526) of the susbtrate molecule facing to the αD (residues 349-354) and αF-αG loop (residues 460-471) of the enzyme acting molecule with a total surface area of ∼1900 Å^2^. A closed up of the interface showed that two acidic residues D521 and E520 (αI) form a salt-bridge with R463 (αF-αG loop). D521 main chain oxygen makes polar contacts with T524 and S525 (αI′) main chain nitrogen atoms. T526 and S525 (αI′) coordinated with K354 (αD) via weak hydrogen bonding (4 Å). This network of hydrogen bonds and electrostatic interactions was further complemented by the coordination of Q278 and G279 (P-loop) main chain atoms in the enzyme acting molecule with the side chain of Q529 at the c-terminal on the susbtrate kinase (Fig. 6B). In this scenario, both molecules adopted a disposition such that we can visualize the c-Src c-terminal Tyr 530 of the susbtrate molecule positioned for ready entry into the active site of the enzyme to undergo intermolecular phosphorylation. This disposition is analogous to the one observed in the crystal structure of the Csk-c-Src complex where the interface was formed by the c-terminal αI helix with the preceding αH-αI loop of c-Src and the αD helix of Csk at the entrance of the active site (Levinson et al., 2008). Superimposition with structures of protein kinases in complex with different peptides substrates: PKA (PDB 1ATP), PKB (PDB 1O6K, 1O6L), PhK (PDB 2PHK) and IRK (PDB 1RK3) revealed that tyrosine and serine/threonine specific substrates follow different paths along the susbtrate binding cleft, and that the c-terminal fragment of c-Src including Tyr530 could readily accommodate onto those paths being compatible with susbtrate binding and phosphorylation (Fig. 6C). To confirm that Tyr 530 auto-phosphorylation was driven by an intermolecular component we performed time course auto-phosphorylation assays at increasing enzyme concentrations (from 0.25 to 2 μM) (Fig. 6D). We observed a concentration dependent effect, as indicated by WB using a phospho-specific antibody. Together our structural and biochemical data demonstrate that the c-terminal Tyr 530 is a de facto c-Src auto-phosphorylation site with a strong intermolecular component.

**Figure 6.**
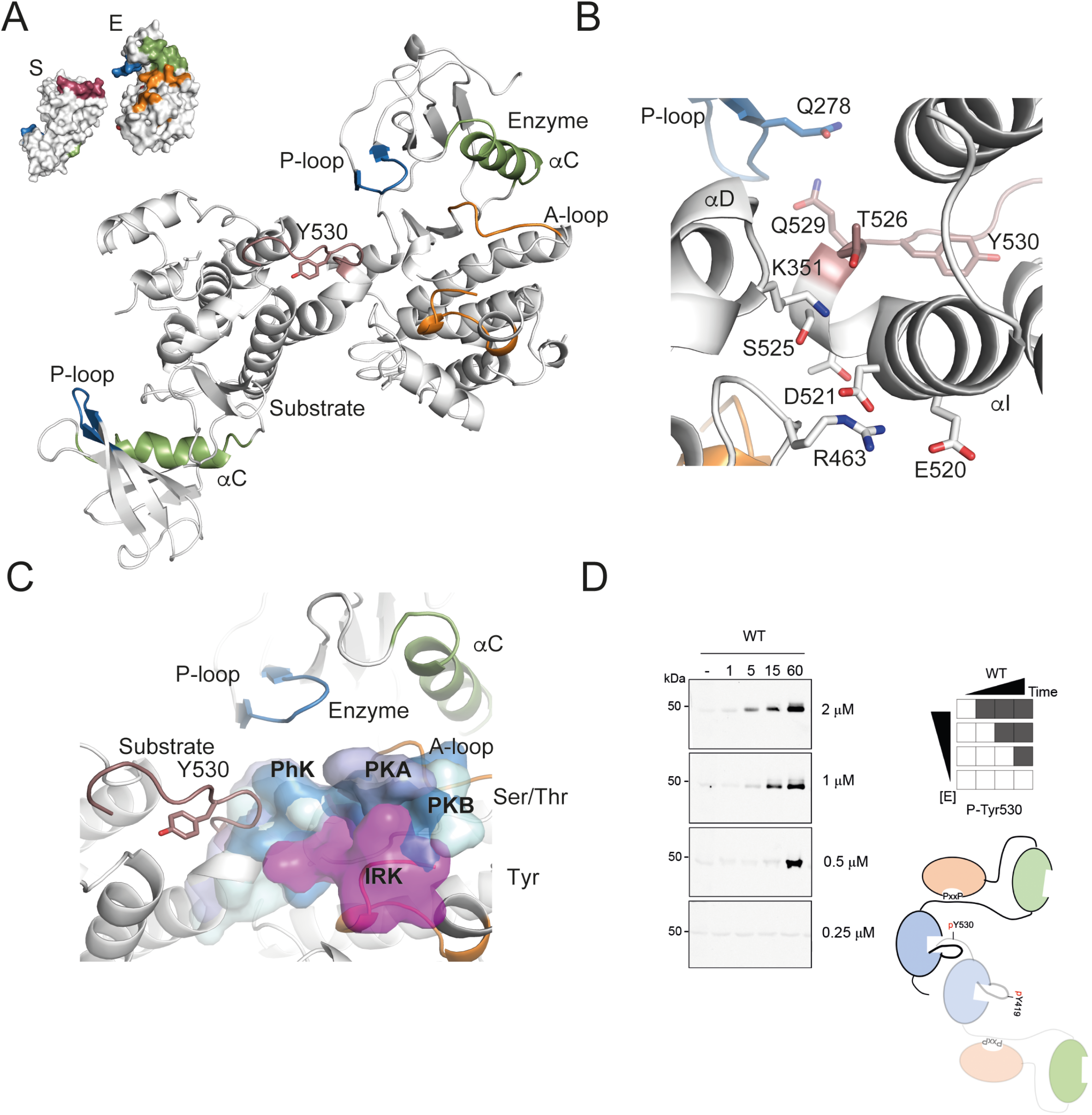
A crystallographic snapshot of c-terminal Tyr530 intermolecular phosphorylation. (A) Cartoon representation of the two molecules (chain A and B) of the asymmetric unit of the c-Src KD-Ponatinib crystal structure where the back side of the c-lobe of one protomer faced the susbtrate binding area located at the base of the active site cleft of the other. Selected residues and secondary structure elements are color-coded as in figure 4. (B) Close-up view of the interface between enzyme and susbtrate molecules showing secondary structural-elements and interacting residues from A. (C) Superimposition of c-Src KD-Ponatinib crystal structure, with previously described structures of protein kinases in complex with different peptides substrates: PKA (PDB 1ATP, light blue), PKB (PDB 1O6K blue, 1O6L cyan), PhK (PDB 2PHK, soft grey) and IRK (PDB 1RK3, magenta) in surface semitransparent representation. (D) WB analyses of an in vitro time-course auto-phosphorylation assay with a c-Src WT 3D-increasing concentrations (from 0.25 to 2 μM) of c-Src WT (3D construct) in the presence of saturating concentrations of ATP (1 mM) and MgCl_2_ (2 mM) for 0–60 min. Total c-Src protein was evaluated by Coomassie staining.

### 9. A c-terminal palindromic phospho-motif controls allosterically c-Src function

We hypothesized that c-terminal truncated viral- and colorectal cancer (CRC)-associated variants may impact on the identified phospho-switch (Fig 7A). To test this premise, we use in time course auto-phosphorylation assays a v-Src 3D surrogate construct lacking the complete c-terminal segment from residue 528 to 536 (Hunter and Sefton 1980) and a colorectal cancer CRC-truncated mutant at codon 531 missing aa 531-536 (Irby et al 1999) (Fig. 7A). Out of the two truncated forms, the c-terminal mutant at codon 531 have an apparent detrimental effect on the catalytic activity as indicated by WBs using a total phospho-tyrosine antibody (Fig. 7B). Unexpectedly, the c-Src 531X mutant, despite having a phosphorylable c-terminal Tyr 530, lacked also the capacity to auto-phosphorylate on this residue, meaning that residues 531 to 536 are required for c-terminal auto-phosphorylation. These data however did not recapitulate the catalytic rates for ATP observed by these variants (Fig. 7C), which was indicative of an otherwise intact catalytic activity. In the c-Src crystal structure we identified a palindromic motif flanking the c-terminal Tyr 530 on the c-tail (PQYQP, aa 528-532) that is disrupted by the c-terminal truncations (Fig. 7D and Fig. 6 A-B). In particular, the side chain of Q529 from the c-tail of the susbtrate molecule is coordinated by hydrogen bonding with Q278 and G279 main chain oxygens (P-loop) of the enzyme acting molecule (Fig. 7D and 6B). These intermolecular contacts were complemented by an intramolecular network of interacting residues, where the c-terminal N535 (c-tail) main chain nitrogen coordinated with side chain of E489 (αG-αH loop) and the side chain of R362 (αE) formed polar contacts with main chain oxygen of G533 (Fig. 7D). These interactions suggested that a perturbation of the palindromic phospho-motif by c-terminal cancer associated variants could affect the kinase-susbtrate interface and hence disturb the phosphorylation switch. In order to evaluate the catalytic function of the c-terminal variants we used intact susbtrate surrogates (i.e. KD K298M constructs) in time-course phosphorylation experiments (Fig. 7E). A quick inspection of the results reveals that both c-terminal variants display comparable catalytic activities to the wild-type as indicated by total-phosphorylation levels of the susbtrate (Fig. 7E). Next, we explored the non-catalytic properties of *de facto* susbtrates (Fig. 7F) in phosphorylation experiments using 3D-K298M constructs with and without c-terminal residues 528-536 (528X) or 531-536 (531X) versus an active c-Src kinase. Surprisingly, both c-terminal variants were also equally and efficiently phosphorylated, which indicate a lack of effect on the non-catalytic properties as a susbtrate. However, and most strikingly, they result in the allosteric inhibition of c-Src Tyr 530 and Tyr 419 auto-phosphorylation on the active kinase (Fig. 7F). We noticed that the allosteric impediment on Tyr 419 auto-phosphorylation was rapid and transient, and recovered over time (Fig. 7F). On the contrary, the inhibitory effect on the c-terminal Tyr530 phosphorylation was more sustained and significant over time. These data provide evidence of the existence of a complex allosteric node at the c-terminus of c-Src controlling the crosstalk between susbtrate- and enzyme-acting kinases, and how perturbation of this allosteric phospho-switch drive c-Src dysfunction in cancer.

**Figure 7.**
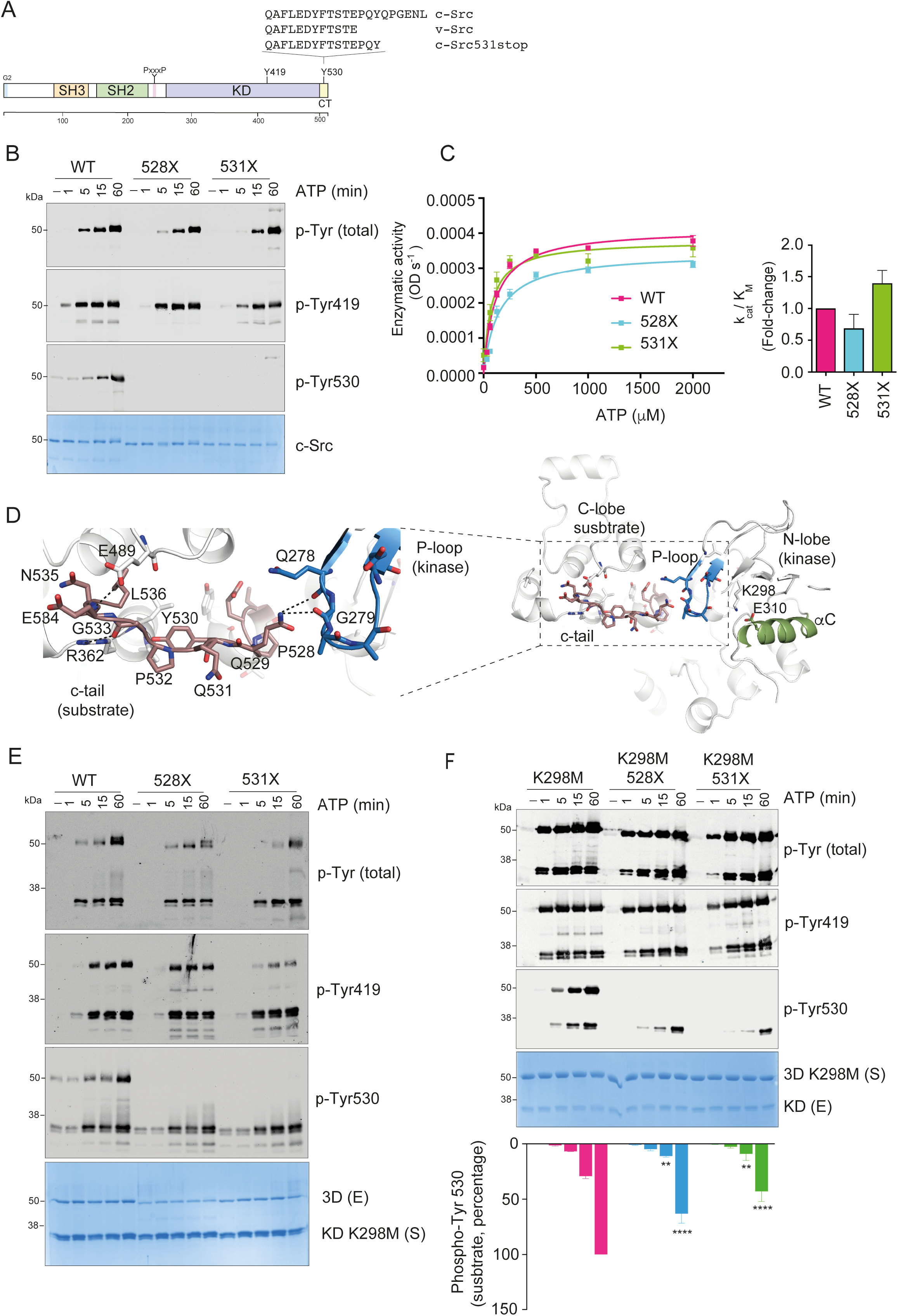
A c-terminal palindromic phospho-motif drives c-Src disfunction in cancer. (A) Schematic diagram of the functional domains and main auto-phosphorylation sites of c-Src. Different c-terminal sequence variants are depicted. (B) WB of samples from a time-course auto-phosphorylation experiment with c-Src WT, v-Src and 531X (3D-construct, 1 μM) in the presence of ATP (1 mM) and MgCl_2_ (2 mM) for 0–60 min using the indicated antibodies. Total amount of protein was visualized by Coomassie staining. Phospho-tyrosine 216 quantification signal, data represent mean ± SEM of 3 experiments (n=3), * p < 0.05, one-way ANOVA test. (C) Enzyme kinetics and catalytic efficiency constant (k_cat_/K_M_, fold-change) for ATP using Src WT, v-Src and 531X (3D-construct, 1 μM final concentration) at fixed concentration (2-4 mg/ml) of Abl peptide. Data represent the mean ± SEM, of 3 experiments in duplicate (n=3). (D) Cartoon representation of the kinase-susbtrate (chain A and B) engagement of two c-Src molecules in the crystal structure. Left panel, close in view of the c-terminal aa sequence of c-Src containing the palindromic PQYQP motif at the interface between the two molecules showing residues coordinated by intra- and inter-molecular interactions. (E) WB of samples from a time-course phosphorylation experiment with c-Src WT, v-Src and 531X (3D-construct, 1 μM) in the presence of a c-Src KD K298M (3 μM) for 0–60 min using the indicated antibodies. Total amount of protein was visualized by Coomassie staining. (F) WB of samples from a time-course phosphorylation experiment with c-Src KD WT (1 μM), in the presence of susbtrate surrogates c-Src, v-Src and 531X K298M (3D-construct, 3 μM) for 0–60 min using the indicated antibodies. Total amount of protein was visualized by Coomassie staining. Phospho-tyrosine 419 and 530 quantification signal on the active kinase molecule, data represent mean ± SEM of 3 experiments (n=3), *** p = 0.0005, **** p = 0.0001, 2-way ANOVA test.

## DISCUSSION

In this study we uncover an allosteric mechanism controlling c-Src intrinsic activity by auto-phosphorylation. Our work dissects a sequential and coordinated cis-to-trans phosphorylation switch connecting the activation loop and c-terminal segment that is required for enzyme specificity and non-catalytic function as a susbtrate. According with this, our crystal structure captures the arrangement of the susbtrate and active kinases during inter-molecular auto-phosphorylation. Furthermore, we identify a c-terminal palindromic phospho-motif containing Tyr 530 on the susbtrate molecule engaging the P-loop of the active kinase. Functional evaluation of cancer related c-terminal variants targeting the phospho-motif provides evidence of the existence of a complex allosteric node at the c-terminus of c-Src modulating the crosstalk between susbtrate- and enzyme-acting kinases.

First of all, we applied a tripartite expression system for heterologous production of c-Src in bacteria (Fig. 1). This system allowed us to increase protein solubility 5-to-10-fold compared with previous bi-partite expression systems where c-Src was co-expressed with YopH or GroEL (Seelinger et al. 2005 and Tong et al 2017). By applying a systematic biochemical and mass spectrometry approach we identify Tyr 530 at the c-terminal segment as a de facto c-Src auto-phosphorylation site with slow time resolution kinetics and strong intermolecular component (Fig. 2B-D and Fig. 7D). These data are further supported by a crystal structure capturing a snapshot of the c-terminal segment of a c-Src susbtrate molecule positioned for ready entry into the active site of the kinase acting molecule (Fig. 7A-C). These data together with: i) results demonstrating that the intrinsically disordered N-terminal region of c-Src does not promote direct dimerization in either the apo nor the ATP-complexed states (Fig. S1A and B), and ii) the lack of binding effect seen on both thermal stability and catalytic activity in experiments using myristoylated G2-derived peptides (Fig. S1 C and D), made us to conclude that contrary to the c-Abl paradigm (Hantschel et al., 2003; Nagar et al., 2003), c-Src myristoylation on G2 play a compartmentalization rather that a functional or catalytic role. Second, we dissected three sequential and coordinated steps in the mechanism of c-Src auto-phosphorylation that couple each phosphorylation event to a specific conformational and functional state (Fig. 2 and 3). In an initial step, a c-Src molecule adopts predominantly a closed inactive configuration with an αC-out and the catalytic salt-bridge (catalytic tether) between K298 and E310 substituted by the E310-R422 tether, based on our SAXS data (Fig. S2) and the crystal structure PDB: 2SRC. It is likely that a partially extended configuration with a flexible and dynamic activation loop can be adopted as a transition to the active state based on PDB: 1Y57. Auto-phosphorylation on activation loop Tyr 419 takes places rapidly in cis-as a priming step in the absence of c-terminal phosphorylation, very likely in an open conformation where the SH2 domain is not engaged with c-terminal Tyr 530 (see graphic abstract). This step requires the transition of the αC helix from the outer to the inner state, so coordination of residues required for catalysis can take place. This priming step switches then to an inter-molecular phosphorylation at later stages compatible with the release and extension of the activation loop to be exposed and accessible to another kinase molecule (step 2). This is followed by the inter-molecular phosphorylation of the c-terminal Tyr 530 by another c-Src kinase molecule (step 3). The first step is consistent with a lack of enzyme concentration effect seen at early time points in the auto-phosphorylation reaction for Tyr 419 (Fig. 4A) and a crystal structure of c-Src (PDB: 2SRC), in which Tyr 419 is pointing to the active site and coordinated with the HRD motif (4 Å to R385) at the same time that the γ-phosphate of the ATP analog coordinates with the DFG motif (4.8 Å from D407), and in close range of Tyr 419 (Xu et al., 1997).The second step is consistent with an enzyme concentration effect seen at later time points in the Tyr 419 auto-phosphorylation reaction (Fig. 4A) and a crystal structure of a phosphorylated c-Src KD in active conformation with an extended activation loop (PDB: 1YI6) positioned readily for intermolecular phosphorylation by another kinase. The third step is supported by our structural and biochemical data demonstrating that Tyr 530 is a de facto c-Src autophosphorylation site (Fig. 2B and D) with a strong intermolecular component driven by a protein concentration effect seen during the auto-phosphorylation reaction (Fig. 6D). This final step is fully dependent on the previous phosphorylation of Tyr 419 on the activation loop (see figure 3A and B and 4B), so phosphorylation of both activation loop and c-terminal segments phosphorylation are sequential and coordinated events (see graphic abstract and Fig. S6). Strikingly, perturbation of such sequential and coordinated mechanism by a c-terminal Y530F mutant bypasses the intermolecular phosphorylation of the activation loop, so it displays a unique cis-component for the mechanism of Tyr 419 auto-phosphorylation (Fig. 4A).

Third, our data using catalytically dead c-Src K298M variants as intact susbtrate surrogates provide solid evidence about how “activating” Tyr419 plays a key non-catalytic role in the presentation of the activation loop of c-Src as a susbtrate to another kinase molecule (Fig. 4C). On the catalytic facet, rather than having a significant impact on the overall c-Src enzymatic activity (Fig. 3B and 4B), activation loop Tyr419 dictates enzyme specificity towards substrates (Fig. 3B and C). These data are consistent with the partial phenotypic rescue observed in different tissues by Drosophila Src42A (Fig. 5A-D) lacking the activation loop tyrosine (Y400F). Differences in the strength of the phenotypes observed when compared with the expression of the WT allele could be explained by i) mutant protein is less efficient functionally, i.e. is lacking the non-catalytic facet and/or ii) differences in susbtrate specificity related to different tissues types. In the case of tracheal tissue (Fig. 5C and D), it is plausible that in a wild-type background, the lack of susbtrate presentation properties by the activation loop mutant has no dominant effect and is compensated by the wild-type allele, which is active both as an enzyme and susbtrate. On the contrary, in a loss of function genetic background using the SrcF80 allele, whose product lacks enzymatic activity, but retains susbtrate function, the activation loop mutant allele has a dominant effect by rescuing the activity of SrcF80 in trans via phosphorylation. Furthermore, by using also K298M susbtrates surrogates, we uncover an allosteric node at the c-terminal palindromic phospho-motif. This allosteric switch was demonstrated by the fact that contrary to wild-type or a Y530F mutant, a c-terminal oncogenic variant without residues 531 to 536 have a detrimental effect on the phospho-tyrosine activity and is unable to auto-phosphorylate on Tyr 530 (Fig. 7B). These data do not match the catalytic activity observed by these variants using both peptides and intact K298M susbtrate surrogates when they are compared with the wild-type (Fig. 7C and E). Furthermore, these variants are efficiently phosphorylated as intact susbtrate surrogates by an active kinase molecule, however and most strikingly the susbtrate molecules with c-terminal variants have a detrimental allosteric effect on the activity of the active molecule (Fig. 7F). The difference in the degree of the allosteric effect observed between the 528X and 531X constructs are not clear to us. One may speculate that residues Q531 and P532 could have an activating input on the phosphorylation of Tyr 530, whereas residues P528 and Q529 have a repressive role via specific contacts between the c-terminal of the susbtrate molecule and the p-loop of the kinase.

Pharmacological inhibition of driver protein kinases by small molecule inhibitors is one of the main therapeutic strategies to treat cancer patients. We solved the crystal structure of c-Src KD in complex with Ponatinib (Fig. S5A), a type-II tyrosine kinase inhibitor used for the treatment of chronic myeloid leukemia (CML) and Philadelphia chromosome–positive (Ph+) acute lymphoblastic leukemia (ALL) (O’Hare et al., 2009; Zhou et al., 2011). This crystal structure resembles to some extent the previously solved c-Src KD-Imatinib crystal structure, and despite sharing the same compound binding pose and DFG-out conformation (Fig. S5A and B) there were significant differences in the arrangement of important secondary structural elements like the P-loop, αC helix and the αC-β3 loop (Fig. S5F). Superimposition with a c-Src KD apo structure (PDB 2OIQ, chain B), which adopted a DFG-in active conformation, revealed a high degree of analogy on that particular region compared with our c-Src-Ponatinib crystal structure. Specifically, in both structures the P-loop was folded over the active site resembling also the configuration seen in the c-Abl KD-Imatinib complex (Schindler et al., 2000). This configuration provides a water-excluding enclosure for the drug increasing the affinity for the ligand and the nucleotide for catalysis. In addition, both the αC helix the β3-αC loop were adopting a lower position compared with the c-Src-Imatinib DFG-out crystal structure (Fig. S5F). While these are all structural features seen in active-like kinases, these inhibitors present in the crystal structures cannot be classified as active per se because the disruption of the linear architecture of the R-spine. This has important implications for the susceptibility to gate keeper resistant mutations. A more compacted αC helix towards the active site (i.e. αC-in) restricts the druggable space as indicated by the distance between gate keeper residue T341 and the catalytic lysine K298 (8.0 Å versus10.4 Å) and as a consequence is more susceptible to resistance by gate-keeper mutations.

Our work provides answers to two important questions for a better understanding of the role of c-Src in human cancers. On one hand, hyper-activation of c-Src in human cancers might result from the perturbation of intrinsic mechanisms of auto-phosphorylation and allosteric control. Large-scale genomic sequencing projects indicate that gene amplification and activating mutations in c-Src do not play a significant role in human tumor biology (Bailey et al., 2018; Curtis et al., 2012). In fact, paradoxical high levels of phosphorylated c-Src in both activation and c-terminal segments are found in aggressive cancer types such TNBC (Nelson et al., 2020) and NSCLC (unpublished). We have data supporting these findings where a previously phosphorylated c-Src protein (90-120 min) is active and able to phosphorylate an intact susbtrate surrogate with faster kinetics than the non-phosphorylated protein (Fig. S7). On the other hand, c-Src may function independent of CSK, which may not always play an anti-oncogenic role through the negative regulation of c-Src in carcinogenesis. In this line, elevated expression of CSK in human cancer cell lines appears to correspond to elevated c-Src protein-tyrosine kinase activity (Watanabe et al., 1995) e.g. CSK is widely expressed in HT-29 and SW620 cells that contain high-levels of active c-Src (Li et al., 1996). Furthermore, genetically engineered mouse models showed that in the absence of CSK, c-Src phosphorylation at Tyr 527 in vivo was reduced to about 20-50% of the level in wild-type cells. These results additionally supported the notion that auto-phosphorylation or another kinase is also involved in c-terminal tyrosine phosphorylation (Bagrodia et al., 1991; Imamoto and Soriano, 1993; MacAuley et al., 1993). In fact, our studies identifying a sequential and coordinated phospho-switch that links the activation and c-terminal segments supports and reinforces this hypothesis.

Altogether, we dissect a phosphorylation switch connecting the activation and c-terminal segments that controls c-Src catalytic and non-catalytic functions. We visualized by X-ray crystallography the intermolecular phosphorylation mechanism between enzyme and susbtrate kinases and identify a functionally relevant c-terminal palindromic phospho-motif on the susbtrate molecule that interacts with the P-loop of the kinase acting molecule. Evaluation of cancer related c-terminal variants perturbing the phospho-motif provide evidence of the existence of a complex allosteric node at the c-terminus of c-Src controlling the crosstalk between susbtrate- and enzyme-acting kinases, and how perturbation of this allosteric phospho-switch drive c-Src dysfunction in cancer. The molecular switches here described are de facto “vulnerabilities” that can be therapeutically exploited for the development of next generation c-Src inhibitors e.g. targeting non-catalytic and/or allosteric features.

## Graphical Abstract

Proposed model for the regulation of c-Src function and conformational landscape by intrinsic auto-phosphorylation. First, a c-Src molecule adopts predominantly a closed **inactive** conformation with an αC-out and the catalytic salt-bridge (catalytic tether) between K298 and E313 substituted by the E313-R422 tether, based on our SAXS data (Fig. S2) and the crystal structure PDB: 2SRC. It is likely that a partially extended configuration with a flexible and dynamic activation loop can be adopted as a transition to the active state based on PDB: 1Y57. Upon ATP and susbtrate accessibility the αC moves to an in-configuration and the catalytic tether (K298-E310) is restored for fast intramolecular phosphorylation on activation-loop Tyr 419 that results in an extended and exposed activation loop based on crystal structure PDB:1YI6. This step is followed by an intermolecular phosphorylation switch on Tyr 419. Next, c-terminal Tyr 530 from the susbtrate acting molecule is positioned for ready entry and phosphorylation by an enzyme acting molecules based on our crystal structure PDB: 7OTE. Finally, a double phosphorylated c-Src adopts a close but active conformation based on our data (Fig. S7) This model has important implications in cancer and provides a plausible molecular explanation for: i) the high levels double-phosphorylated c-Src on both activation and c-terminal segment tyrosines found in aggressive types of breast cancer such TNBCs, and ii) the perturbation of the allosteric phospho-switch by c-terminal truncated cancer variants. Non-phosphorylable c-terminal variants adopt predominantly a partially extended configuration as seen in PDB: 1Y57 and undergo intramolecular phosphorylation on the activation loop. Intermolecular phosphorylation of the A-loop is favored by intermolecular contacts between the P-loop of the enzyme kinase and the c-terminal of the substrate molecule (e.g. Q278-Q531) that in the case of these variants are missing. C-terminal intermolecular phosphorylation is abolished in this setting due to the absence of the phospho-motif and the lack of contacts between the P-loop of the active kinase and the c-terminal of the susbtrate molecule (Q278-Q529) as seen in PDB. 7OTE.

## Author contributions

Conception of the study and experimental design (IP-M), manuscript writing, data analysis and figures preparation (IP-M), Drosophila work, data analysis and figure preparation (MLL), mass spectrometry including data analyses (JS and JM), SAXS studies including data analyses and figure preparation (IM), experimental work (NC, JC, PS), X-ray crystallography (NC, PS) and funding acquisition (IP-M).

## Acknowledgements

We are grateful to CNIO for supporting this study, to Clara Santiveri and Ramón Campos from the Spectroscopy and Nuclear Magnetic Resonance Unit (CNIO) for excellent technical support, the Genomics Unit (CNIO), and members from the Kinases, Protein Phosphorylation and Cancer Group for their technical assistance, specially to Moustafa A. Shehata for helping with supplemental figure 6 preparation and Julio Martínez-Torres for helping with recombinant protein expression and cloning the pUAST-Src42A construct. To Eduardo Zarzuela from the Proteomics Unit (CNIO) for assistance with Mass Spectrometry studies. We thank to Nabil Djouder and Javier Klett for helpful comments and advice on the manuscript. The authors would like to thank Diamond Light Source for beamtime (beamline B21, proposal mx30297) and the ALBA Synchrotron Light Facility (XALOC-BL13 beamtime, proposal 2020094499) and their staff for assistance during data collection.

## Materials, data availability and correspondence

The crystallographic coordinates and structure factors for the crystal structure of human c-Src KD in complex with Ponatinib reported in this paper were deposited in the Protein Data Bank (PDD) with the PDB code 7OTE.

