## Supplemental information and figures for "An allosteric switch between the activation loop and a c-terminal palindromic phospho-motif controls c-Src function"

**Supplemental Information *Cuesta et al.***

**1. Extended material and methods**

**2. Supplemental figures**

**Figure S1**

(A-B) Drosophila Src42A purification and MS identification of oligomeric species

(C-D) Evaluation of Myr-G2 versus G2 peptides on binding and activity assays

**Figure S2.** SAXs analysis of un-phosphorylated c-Src 3D construct

**Figure S3.** MS data of c-Src WT auto-phosphorylation

**Figure S4.** MS data of c-Src WT, Y419F and Y530F auto-phosphorylation

**Figure S5.** Enzymatic activity ($K_M$ ATP) at increasing enzyme concentration of c-Src WT, versus Y419F and Y530F mutants

**Figure S6.** Crystal structure of c-Src KD in complex with Ponatinib

**Figure S7.** Functional evaluation of phosphorylated c-Src

**3. Supplemental tables**

**Table S1.** X-ray data quality indicators and statistics

**Table S2.** SAXs data collection and structural and mass parameter

**4. References**

### 1. EXTENDED MATERIALS AND METHODS

#### Plasmids

A pET28a-TEV plasmid codifying an N-Terminal 6xHis tag followed by a Tobacco etch virus (TEV) protease recognition site was used to express human c-Src (UniProtKB P12931) three domain (3D, SH3-SH2-KD aa 84-536) or kinase domain (KD, aa 254-536) codifying sequences. Both pFCDUET-YopH phosphatase and pGKJE8-GroEl/GroEs chaperones plasmids (Takara chaperone plasmid set cat. # 3340) were used for co-expression experiments. These constructs were a kind gift of Dr. Daniel Lietha (CIB, Madrid). For expression of recombinant drosophila c-Src isoform 42A (Src42A, UniProtKB Q9V9J3) we used a pET-His10-Src42A plasmid (Addgene #126674) codifying a full-length drosophila isoform (aa 1-517) with a N-terminal 10x His tag and a Thrombin protease recognition site. For experiments in Drosophila a pUAST-attB plasmid encoding drosophila Src homolog isoform A (Src42A) was cloned by amplifying dSrc42a from a pGEX-Src42a donor plasmid (Addgene #126673) using primers with EcoRI (forward 5´- CTGAATAGGGAATTGGGAATTCATGGGTAACTGCCTCACC-3´) and XbaI (reverse 5´- CCTTCACAAAGATCCTCTAGATCAGTAGGCCTGCGCCTC-3´) restriction sites.

#### Site-directed mutagenesis

Site-directed mutagenesis was performed in order to generate all the point mutants and variants described in this study using the Q5-site directed mutagenesis kit (New England Biolabs) following manufacturer instructions and the indicated primers (see extended material and methods).

K298M-forward 5´- GGTGGCCATCATGACCCTGAAGC - 3´

K298M-reverse 5´- CTGGTGGTACCGTTCCAG - 3´

Y419F-forward 5´- AGACAATGAGGAGACGGCGCGGC - 3´

Y419F-reverse 5´- TCAATGAGCCGAGCCAGC - 3´

Y530F-forward 5´- CGAGCCCCAGTTCCAGCCCGGGG - 3´

Y530F-reverse 5´- GTGGACGTGAAGTAGTCCTCCAGGAAGG - 3´

dY400F-forward 5´- GGAGGACGAATTCGAGGCGCGGG - 3´

dY400F-reverse 5´- TTGATGAGCCTAGCTAAACCAAAGTCG -3´

P528X/Q529X-forward 5´- GTCCACCGAGTAATAGTACCAGCCCG -3´

P528X/Q529X-reverse 5´- GTGAAGTAGTCCTCCAGG -3´

Q531X/P532X-forward 5´- GCCCCAGTACTAGTAGGGGGAGAACC -3´

Q531X/P532X-reverse 5´- TCGGTGGACGTGAAGTAG -3´

Briefly, in an initial exponential amplification step template plasmid DNA (1-25 ng) was incubated with Q5 Hot Start High-Fidelity master mix (2X), 10 µM of forward and reverse primer stock to a final concentration of 0.5 µM and nuclease-free water in a final volume of 12.5 µl. Amplification and cycling conditions were as followed: i) initial denaturation at 98°C 30 sec, ii) 98°C 10 sec, 50–72°C* 30 sec, 72°C 4 min (25 cycles) and iii) final extension 72°C (2 min). For mutagenic primers, we used the $T_a$ provided by the online NEB primer design software, NEBaseChanger™. Next, a kinase-ligase and Dpn-I (KLD) reaction was set at room temperature for 5-30 min, in which a 1 µl of PCR product was incubated with a 2X KLD reaction buffer, 10X KLD enzyme mix and nuclease-free water in a total volume of 5 µl. Q-5 site directed mutagenesis products (5 µl) were transformed in E. coli Q5-DH5α by incubation for 30 minutes on ice and heat shocked at 42ºC for 45 seconds after which samples were placed back on ice for 2-3 minutes prior recovery in 500 µl of antibiotic free LB media at 37ºC and 200 rpms shaking for one hour. Next, 150 µl of recovered culture was plated onto LB-Agar plates with kanamycin (50 mg/ml) and left overnight (o/n) into an incubator at 37 ºC. Next day, one single colony was amplified o/n in 10 ml of LB media with kanamycin (50 μg/mL) and the next morning the bacterial culture was pelleted by centrifugation at 3.000 rpm and plasmid DNA purification was performed using the E.Z.N.A. plasmid DNA mini kit (Omega Bio-Tek Inc) following manufacturer instructions. Briefly, bacterial pellet was resuspended in 250 µL Solution I (resuspension solution) supplemented with RNAse A (1/1000 v/v). Then, the same volume of Solution II (lysis solution) was added, mix and incubated for 2 min. Then, 350 µl solution III (neutralization solution) was added and centrifuged at 13.200 rpms in an eppendorf centrifuge 1545 D for 10 min at room temperature. Supernatant was transferred to a HiBind® DNA mini column and centrifuged for 1 min and flow through discarded. Then, 500 µL HBC solution was added, centrifugated for 1 min and flow through discarded again. Process was repeated with 700 µL of DNA washing buffer. Then, 50 µl of elution buffer was added and centrifuged for another minute on a clean eppendorf. Plasmid concentration and purity was checked by absorbance using a nanodrop (Thermo Scientific NanoDrop One). All mutagenesis products were confirmed by DNA Sanger sequencing at the CNIO Genomics Unit.

**Expression and purification of recombinant proteins**

In order to express high yields of soluble and monodisperse recombinant human c-Src protein we followed a modified protocol (Seeliger et al., 2005) in which in addition to YopH phosphatase, we also co-expressed the chaperone GroEl. For co-expression, pET28a-TEV-hSrc [84-536] and pET28a-TEV-hSrc [254-536] plasmids were co-transformed in E. coli BL21 bacteria strand previously transformed with pFCDUET-YopH and pGKJE8-GroEl/GroEs plasmids and grown in 50 ml of LB media containing kanamycin (50 μg/ml), streptomycin (50 μg/ml) and chloramphenicol (34.5 μg/ml) overnight at 37ºC shaking at 200 rpms in a 250ml Erlenmeyer flask. Next day, the bacterial culture was diluted (1:100) in LB media with the same antibiotic composition and concentration using Fernbach baffled Erlenmeyer flasks. When the bacterial culture reached an optic density (λ 600nm) of 0.15, the culture was cooled to 18ºC and tetracycline was added at a final concentration of 1ng/ml in order to trigger chaperone expression. After approximately one hour and always at an optic density of 0.4-0.5 at λ 600 nm, IPTG was added to a final concentration of 500 μM and culture was left overnight agitating (200 rpms) at 18ºC. To express recombinant dSrc42A we followed the same procedure but with the co-expression of YopH only, so a pET-His10-Src42A plasmid was transformed in BL21 previously transformed with the pCFDUET-YopH construct and tetracycline was not added at any moment.

Next day, bacterial culture was harvested at 3000 rpms and 4ºC with a JLA 8.100 rotor in a Beckman Coulter Avanti J-20XP centrifuge. Supernatant was discarded and pellet was transferred into a 50 ml Falcon centrifuge tube and frozen at -20 ºC for later purification. Alternatively, the pellet was immediately resuspended in 50 ml lysis buffer (50 mM Tris pH 8, 500 mM NaCl, 0.1mM PMSF) and sonicated on ice with a Bioblock Scientific Vibra Cell 75042 sonicator with a model CV33 sound radiator for 4 minutes (9 seconds on-3 seconds off, 37% amplitude). Crude lysate was transferred into a JA 25-50 rotor tubes and centrifugated at 20.000 rpms at 4ºC for 45 min in a Beckman Coulter Avanti J-25 centrifuge. Soluble clarified supernatant underwent a further 10 seconds sonication step and was filtered through a 45 μm Jet Biofilter with a syringe.

Human c-Src was purified by three chromatographic steps (see figure 1). First, lysate was passed through an immobilized metal anion chromatography (IMAC) 5ml column (GE Healthcare HisTrap$^{TM}$ HP) equilibrated with 5 column volumes (CVs) of IMAC buffer A (20mM Tris pH 8, 150mM NaCl, 1mM TCEP, 5% Glycerol) with an GE Healthcare AKTA PURE FPLC at flowrate of 5 ml/min. Next, the column was washed with 90% IMAC buffer A and 10% IMAC buffer B (20 mM Tris pH 8, 150 mM NaCl, 300 mM Imidazole, 1mM TCEP, 5% Glycerol) until absorbance signal at a λ of 280nm was stable (usually 10 CVs), after which a 100% gradient with IMAC buffer B was run in 100 ml (20 CVs) and 20 fractions of 5 ml were collected. Fractions were tested in a 12% SDS-PAGE gel and Coomassie blue staining.

Fractions expressing recombinant c-Src were pooled together and diluted in IEC (ionic exchange chromatography) buffer A (20mM Tris pH 8, 1mM DTT, 5% glycerol) up to 3 times original volume so NaCl concentration was lowered to 50mM. Then, sample was loaded into an IEC column (GE Healthcare HiTrap Q HP column) previously equilibrated with 5 CVs of IEC buffer A at a flowrate of 5ml/min. After loading the column was washed until absorbance signal at λ 280nm was stable (4-5 CVs) and then a 100% gradient with IEC buffer B (20mM Tris pH 8, 500mM NaCl, 1mM DTT, 5% glycerol) was run in 100 ml (20 CVs) and 20 fractions of 5ml each were taken. Fractions were tested in a 12% SDS-PAGE gel and Coomassie blue staining.

Fractions expressing recombinant c-Src were pulled together and mixed with a His-tagged rTEV protease (20-40 μM) in a 1/20 molar TEV/c-Src stoichiometry. Buffer was supplemented with 2mM TCEP and digested at 4ºC o/n. Alternatively, digestion was performed for 2 hours at room temperature. Next, a His-trap reverse step was undertaken to remove the protease and tag of the recombinant protein. Briefly, His-rTEV protease digested c-Src sample (input) was passed through the HisTrap column previously equilibrated with IMAC buffer A at 2 ml/min flowrate with a peristaltic pump (GE Healthcare Pump P-1) and then washed with 5 CVs of buffer IMAC A supplemented with 50 mM imidazole. Flow through (FT) was taken and check together with the input in a 12% SDS-PAGE gel and Coomassie blue staining.

FT was taken and concentrated with a Millipore concentrator with a 10-30 kDa cut-off by centrifugation with an Eppendorf Centrifuge 5810 R at 3.500 rpms at 4 ºC. When c-Src volume was less than 2.5 ml it was injected in a 5ml loop (GE Healthcare) to run a size exclusion chromatography (SEC) with a Superdex 200 16/60 column (GE Healthcare) previously equilibrated with 1.5 CVs of SEC buffer (20 mM Tris pH, 150mM NaCl, 1mM DTT, 5% Glycerol). Fractions of 5 ml were collected and tested in a 12% SDS-PAGE gel and Coomassie blue staining. Protein concentration and purity were checked by absorbance using a nanodrop.

In the case of dSrc 42A purification only two chromatography steps were undertaken: First, clarified lysate was passed through an IMAC 5ml column (GE Healthcare HisTrap HP) equilibrated with 5 CVs of buffer A (20mM Tris pH 8, 150mM NaCl, 1mM TCEP, 5% Glycerol) with an GE Healthcare AKTA PURE FPLC at a flowrate of 5ml/min. After loading the column was washed with 90% buffer A and 10% buffer B (20mM Tris pH 8,150mM NaCl, 500mM Imidazole, 1mM TCEP, 5% Glycerol) until absorbance signal at $\lambda$ 280nm wavelength signal was stable; typically, it was for 7 CVs. Then, a 100% gradient of buffer B was run in 100 ml 20 CVs taking 20 fractions of 5ml each. Fractions were tested in a 12% SDS-PAGE gel and Coomassie blue staining. Fractions expressing recombinant dSrc42A were pulled together and concentrated in a Millipore concentration (30 kDa cut-off) as indicated before. Concentrated sample (2.5 ml) was injected using a 5 ml loop in an AKTA PURE FPLC (GE Healthcare) to run a SEC with a Superdex 200 16/60 column (GE Healthcare). Fractions were tested in a 12% SDS-PAGE gel and Coomassie blue staining. Protein concentration and purity were checked by absorbance using a nanodrop.

**Size-Exclusion Chromatography with Multi-Angle Light Scattering (SEC-MALS)**

A sample volume of 400-500 $\mu$L at a minimum concentration of 100 $\mu$gr was injected in a Superdex 200 Increase 10/300 column (Cytiva) equilibrated in 20 mM Tris pH, 150mM NaCl, 1mM DTT, 5% Glycerol buffer (filtered through a 0.1 $\mu$m filter) and connected to an AKTA Purifier equipment (GE Healthcare). The chromatographic eluent was monitored by three consecutive detectors in series: (1) a multi-wavelength UV-Vis absorbance detector Monitor UV-900 of the AKTA system (GE Healthcare) with a 10 mm path length flow cell, (2) a light scattering DAWN Heleos 8+ (Wyatt Technology) with detectors at eight different angles (from 32 to 141 degrees from the source) using a linearly polarized GaAs laser operating at 665 nm and (3) an Optilab T-rEX (Wyatt Technology) differential refractive index detector with a laser wavelength of 658 nm. The column was equilibrated overnight in running buffer at 0.1 mL/min flow to obtain stable base lines for the detectors before data collection. After that, all the experiments were performed at 0.5 mL/min flow and room temperature (~25 ºC). Before running test samples, a control run with BSA (500 μl at 0.5 mg/ml), a well-characterized monodisperse sample, was carried out to set the alignment and band broadening parameters and the normalization coefficients of the MALS detectors necessary for data analysis. Data collection and analysis were performed using UNICORN 5.10 (GE Healthcare) and ASTRA 6.0.3 (Wyatt Technology) software packages.

**Protein electrophoresis**

Protein electrophoresis were performed with 12% SDS-PAGE gels. The separating gel polymerized with 375 mM Tris pH 8.8, 12% acrylamide/Bis 30% w/v, 0.1% SDS, 0.1% ammonium persulfate (APS) and 0.5% TEMED. Gel staking polymerized with 125 mM Tris-Cl pH 6.8, 12 % acrylamide, SDS 0.1%, APS 0.1% and TEMED 1% (Table 2). These gels were run in electrophoresis buffer (25 mM Tris pH 8.3, 192 mM glycine, 0.1% SDS) using the Mini-PROTEAN system (BIO RAD) at a constant 200mV for 30min using a BIO RAD PowerPac Basic. For Coomassie staining, gel was immersed in Coomassie blue staining solution (10% acetic acid, 40% absolute ethanol and 50% deionized water with 1g/L Brilliant blue 250 R. Once the gel was blue (typically, after 30 min) it was immersed in Coomassie distaining solution (10% acetic acid, 50% absolute ethanol, 40% deionized water).

### Western blotting and antibodies

SDS-PAGE gels were transferred onto nitrocellulose 0.2 μm (Amersham™) or PVDF 0.45 μm (Millipore) membranes using the Mini-PROTEAN system (BIO RAD). PVDF membranes were activated first in ethanol 100% for 10 min. SDS-PAGE gel and membrane were sandwiched between four filter paper slices in mini-PROTEAN system immersed in transfer buffer (25 mM Tris pH 8.8, Glycine 190 mM and ethanol 10%) and run at constant 200 nV for 120 min on ice. Transferred membranes were immersed in blocking solution (10 mM Tris pH 8, 150 mM NaCl, 5% weight/volume (w/v) skimmed powder milk) for 60 min shaking at 15 rpms on a shaker see-saw rocker SSL4 (STUART). After blocking membranes were washed three times with TBS-T (10 mM Tris pH 8, 150 mM NaCl, Tween-20 0.1% v/v) prior incubation with primary antibody solution (TBS-T with BSA 5% w/v) o/n at 4ºC shaking at 15 rpms on a Duomax 1030 (Heidolph) shaker. Antibodies used were phospho-Src Tyr419 (D49G4) CST #6943 and Src (36D10) CST #2109 were diluted at 1/10000 and antibodies phospho-Src Tyr 530 (ThermoFisher) and total phospho-Tyr (p-Tyr-100 CST #9411) at 1:5000. Next day membranes were washed with 20 ml TBS-T 3-to-5 times and immersed in secondary antibody solution (TBS-T skimmed powder milk 5% m/v). Secondary antibodies anti-rabbit or -mouse IgG (DyLight conjugate at 680 or 800 nm (CST #5366 #5151 #5470 #5257) were used at half the dilution factor o the primary for one hour at room temperature protected from light. After incubation with secondary antibodies, membranes were washes 3-to-5 times with TBS-T. Next, membranes were scanned in an Odyssey CLx scanner and images exported.

### In vitro phosphorylation assays

In vitro phosphorylation assays were performed at room temperature using recombinant human c-Src 3D and KD (WT and indicated mutants) and full-length dSrc42A at 1 μM final concentration in buffer (20 mM Tris pH, 150mM NaCl, 1mM DTT, 5% Glycerol, 2mM

$MgCl_2$) and ATP (1 mM) at the indicated time points (unstimulated, 1, 5, 15, 30, 60 and 90 min). Aliquots from the time-course were mixed with 5X Laemmli sample buffer (ThermoFisher) and denaturalized at 95ºC during 1-2 min in a thermoblock AccuBlock (Labnet).

**Enzymatic assays**

Phosphorylation rates of peptide substrates by recombinant c-Src and its variants at 1μM final concentration were determined by using an NADH-coupled pyruvate kinase assay in presence of increasing ATP concentrations. The enzyme-substrate solution (20 mM Tris-Cl, 1 mM $MgSO_4$, 400 mM Phosphoenolpyruvate (PEP), 100 mM NADH, 2450U/ml Pyruvate kinase (PK), 2.26 mg/ml Lactate dehydrogenase (LDH) was prepared at different concentrations of ATP: 0, 0.08, 0.16, 0.32, 0.65, 1.25, 2.5 and 5 mM. The experiments were performed in a 384 well plate (Greiner bio-one) and the NADH consumption was read at 340 nm wavelength during 120 cycles of 30 sec each by using a Victor multilabel plate reader 1420 Multilabel Counter (Perkin Elmer). In order to obtain catalytic rates and kinetic constants by Michaelis-Menten equation, experiments were analyzed using Prism software. The following peptides sequences were used as exogenous substrates: c-Abl (EAIYAAPFAKKK), c-Src Tyr 419 (IEDNEYTARQG), c-Src Tyr 530 (STEPQYQPGEN), c-Src G2 (GSNKSKPKDASQRRR), c-Src Myr (Myr-GSNKSKPKDASQRRR) and RET Tyr 900/905 (DVYEEDSYVK).

**Differential Scanning Fluorometry (DSF)**

To evaluate the thermal stability of recombinant c-Src WT and indicated variants in the absence of (apo) and in complex with Ponatinib, we applied two different scanning fluorometry methods. First, an indirect SYPRO Orange-based method. For this assay the total reaction volume was adjusted to 40 µL at 1-2 µM protein, 10 µM inhibitor, and 2 x SYPRO Orange concentrations subjected to a gradient of temperature from 20 to 95 °C. Fluorescence was measured on an Applied Biosystem 7300 Real-Time PCR system. Second, a direct method based on changes in intrinsic fluorescence upon a quick gradient of temperature was measured using a tycho instrument (Nanotemper) at 1-2 µM protein, 10 µM inhibitor concentration and following manufacturer´s instructions.

**Drosophila strains**

Drosophila melanogaster strains were maintained and raised at 25ºC under standard conditions. The stocks used are the following ones: GMRGal4 (eye expression, BSDC 9146), salEPvGal4 (wing expression, BDSC 80573), btlGal4 (tracheal expression, gift of

S. Hayashi), $src^{F80}$/CyOlacZ; UASSrc/TM6 dfdYFP (gift of S. Luschnig); UASSrcY400F/TM6 dfdYFP (this work).

Balancer chromosomes CyO, TM3 or TM6 marked with LacZ, GFP or YFP were used to follow the mutations and constructs of interest in the different chromosomes. Transgenes were generated by injecting the pUASTattBSrc42aY400F construct to obtain directed insertions at 68E by the “Transgenesis Service” of the “Centro de Biología Molecular Severo Ochoa” (CBM, Madrid). Transgene expression was achieved using the Gal4/UAS system (Brand and Perrimon, 1993) at 25°C (GMRGal4) or 29°C (btlGal4 and salEPvGal4).

**Immunohistochemistry**

Embryos were stained following standard protocols. Embryos were fixed in 4% formaldehyde (Sigma-Aldrich) in PBS1x-Heptane (1:1) for 20 min. Embryos transferred to new tubes were washed in PBT-BSA blocking solution and shaken in a rotator device at room temperature. Embryos were incubated with the primary antibodies in PBT-BSA overnight at 4ºC. Secondary antibodies diluted in PBT-BSA (and for the CBP staining) were added after washing and were incubated at room temperature for 2-5 h in the dark. Embryos were washed, mounted on microscope glass slides with Fluoromount-G (Southern Biotech) and covered with thin glass slides. Primary antibodies used were goat anti-GFP (1:600) from Roche and chicken anti-β-gal (1:200) from Abcam. Alexa Fluor 488, 555, 647 (Invitrogen) secondary antibodies were used at 1:300 in PBT 0.5% BSA. CBP (Chitin Binding Protein, produced by N. Martín in Dr. Casanova's lab, New England Biolabs Protocol) was used as a secondary antibody at 1:300 to detect chitin and visualize the tracheal branches.

### Image acquisition

Images from fixed embryos were taken using Leica TCS-SPE with the 20x and 63x immersion oil (1.40-0.60; Immersol 518F – Zeiss oil) objectives and additional zoom. Settings were adjusted for the different channels prior to image acquisition. Z-stack sections of 0.24-0.5 µm were acquired. The images were imported and processed using Fiji (ImageJ 1.49b) and Photoshop for measurements and adjustments, and assembled into figures using Illustrator.

U.S.A.). The crystallographic coordinates and structure factors for the crystal structure of human c-Src KD in complex with Ponatinib reported in this paper is PDB: 7OTE. For data statistics, see table 1.

**Small angle X-ray scattering (SAXs)**

SAXS experiments were conducted at the beamline B21 of the Diamond Light Source (Didcot, UK) (Cowieson et al., 2020). A sample of 40 ul of Src WT (3D-construct) at concentration of 3 mg/ml was delivered at 20ºC via an in-line Agilent 1200 HPLC system in a Superdex 200 Increase 3.2 column, using a running buffer composed by 20 mM Tris pH 8.0, 150 mM NaCl, 1% glycerol and 1 mM DTT. The continuously eluting samples were exposed for 300s in 10s acquisition blocks using an X-ray wavelength of 1 Å, and a sample to detector (Eiger 4M) distance of 3.7 m. The data covered a momentum transfer range of 0.0032 < q < 0.34 Å-1. The frames recorded immediately before elution of the sample were subtracted from the protein scattering profiles. The Scåtter software package (www.bioisis.net) was used to analyse data, buffer-subtraction, scaling, merging and checking possible radiation damage of the samples. The $R_g$ value was calculated with the Guinier approximation assuming that at very small angles $q < 1.3/R_g$. The particle distance distribution, $D_{max}$, was calculated from the scattering pattern with GNOM, and shape estimation was carried out with DAMMIF/DAMMIN, all these programs included in the ATSAS package (Petoukhov et al., 2012). The proteins molecular mass was estimated with GNOM. Interactively generated PDB-based homology models were made using the program COOT (Reference CCP4) by manually adjusting the X-ray structures obtained in this work, into the envelope given by SAXS until a good correlation between the real-space scattering profile calculated for the homology model matched the experimental scattering data. This was computed with the program FoXS (Schneidman-Duhovny et al., 2016).

### 2. SUPPLEMENTAL FIGURES

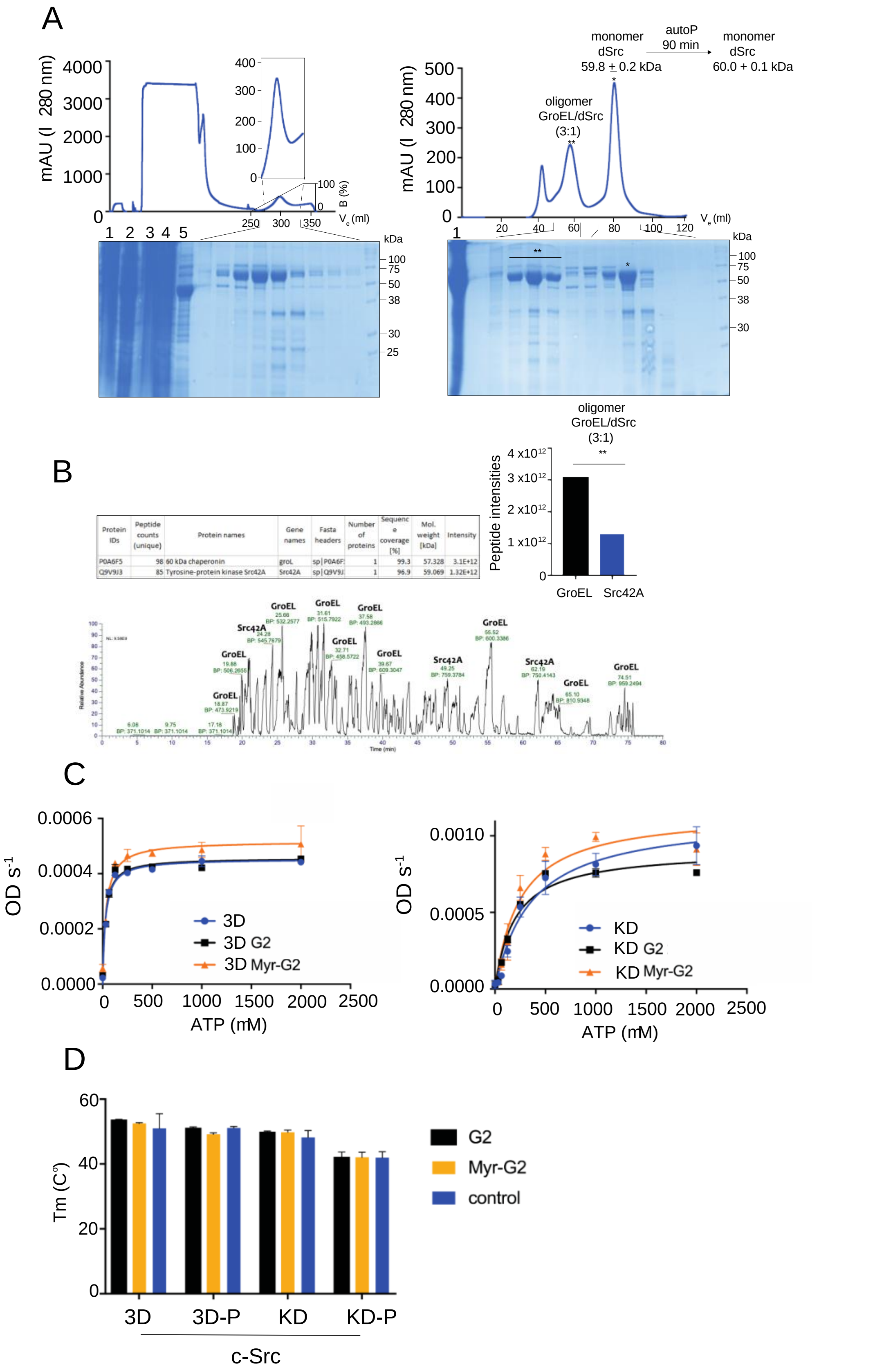

| Protein IDs | Peptide counts (unique) | Protein names | Gene names | Fasta headers | Number of proteins | Sequence coverage [%] | Mol. weight [kDa] | Intensity |
|---|---|---|---|---|---|---|---|---|
| P0A6F5 | 98 | 60 kDa chaperonin | groL | sp\|P0A6F | 1 | 99.3 | 57.328 | 3.1E+12 |
| Q9V9J3 | 85 | Tyrosine-protein kinase Src42A | Src42A | sp\|Q9V9J | 1 | 96.9 | 59.069 | 1.32E+12 |



**Figure S1.** Purification and characterization of Src42A species by SEC-MALS and MS and functional characterization of Myr-G2 derived peptides. (A) IMAC chromatogram using a HisTrap column (5 ml). Indicated fractions were run on an SDS-PAGE and stained with Coomassie: 1 crude lysate, 2 insoluble fraction, 3 clear lysate, 4 wash, 5 flow-through, left panel. SEC chromatogram using a Superdex 200 16/60 column. Indicated fractions were run on an SDS-PAGE and stained with Coomassie, right panel. (C) Base peak chromatogram (BPC) from LC-MSA/MS analysis showing stochiometric relationship of GroEL and dSrc (Src42A) in the oligomeric fraction obtained from the SEC. (D) Enzymatic assays performed with c-Src 3D or KD (both at 1 $\mu$M) in apo estate (0-P) or phosphorylated (P, 90 min) incubated for 45 min with a 2.5 molar excess of G2 (GSNKSKPKDASQRRR) and Myr-G2 (Myr-GSNKSKPKDASQRRR) peptides using c-Abl peptide (4mg/ml) as a phosphorylable susbtrate. DSF data represented in a column plot showing the melting temperature of c-Src 3D and KD unphosphorylated (0-P) and phosphorylated c-Src (P, 90 min) species in apo (control), Myr-G2 and G2 pre-incubated samples. Data represented are the mean ± SEM of 2 experiments.

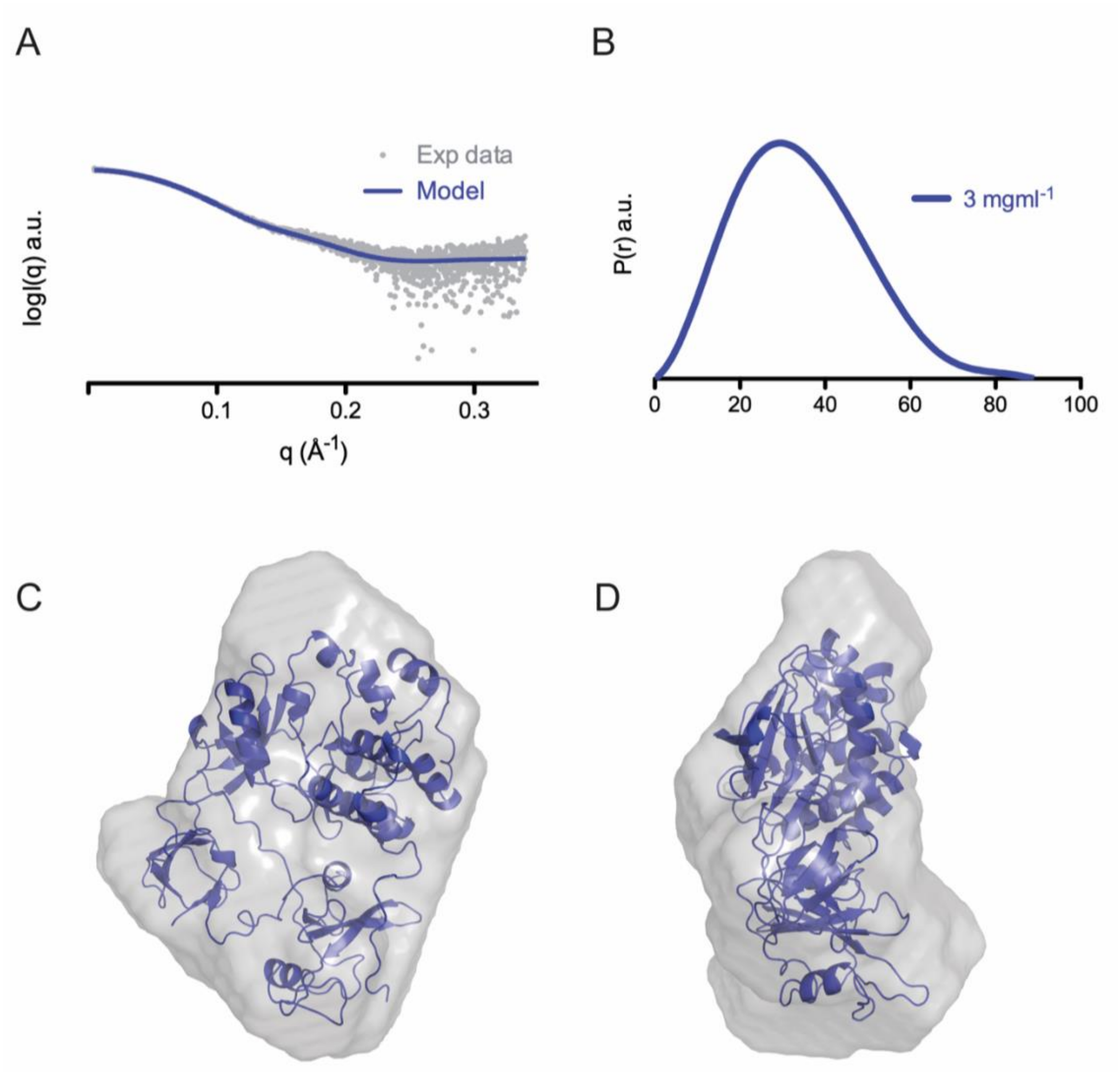


**Figure S2.** Analysis by SAXS of Src WT domain organization in solution

(A) Experimental scattering curve (grey dots) and theoretical scattering curve, in blue, computed for the model (smooth), where Src WT adopts a closed conformation as found in the PDB: 2SRC.

(B) Normalized pair-distance distribution function P(r) for Src WT (blue graph). The data were offset vertically for clarity, a.u., arbitrary units.

(C) Overlaying of the ab initio determined SAXS envelope for Src-WT (in pale grey), with the model based on the crystal structure (PDB: 2SRC), in blue.

GVTTFVALY$_{93}$DY$_{95}$ESRTETDLSFKKGERLQIVNNT$_{117}$EGDWWLAHSLS$_{128}$T$_{129}$GQT$_{132}$
GY$_{139}$IPSNY$_{139}$VAPSDSIQAEEWY$_{152}$FGKITRRESERLLLNAENPRGT$_{174}$FLVRESET$_{182}$T
KGAY$_{187}$CLSVSDFDNAKGLNVKHYKIRKLDSGGFY$_{216}$ITSRTQFNSLQQLVAY$_{232}$Y$_{233}$SK
HADGLCHRLTTVCPTSKPQTQGLAKDAWEIPRESLRLEVKLGQGCFGEVWMGT$_{288}$W
NGTTRVAIKTLKPGTMSPEAFLQEAQVMKKLRHEKLVQLYAVVSEEPIY$_{338}$IVTEY$_{343}$M
SKGSLLDFLKGETGKYLRLPQLVDMAAQIASGMAY$_{379}$VERMNY$_{385}$VHRDLRAANILVG
ENLVCKVADFGLARLIEDNEY$_{419}$TARQGAKFPIKWTAPEAALY$_{439}$GRFTIKSDVWSFGIL
LTELTTKGRVPY$_{466}$PGMVNREVLDQVERGY$_{482}$RMPCPPECPESLHDLMCQCWRKEP
EERPTFEYLQAFLEDYFTSTEPQY$_{530}$QPGENL

| Residue | Described at biochemical and/or cellular level | Auto-phosphorylation or phospho-site for other kinases | Ref. |
|---|---|---|---|
| Y93 | Yes | Phospho-site | [1] |
| Y95 | No | - | |
| T117 | No | - | |
| S128 | No | - | |
| T129 | No | - | |
| T132 | No | - | |
| Y134 | Yes (mouse) | Phospho-site | [2] |
| Y139 | Yes (mouse) | Phospho-site | [2], [3] |
| Y152 | No | - | |
| T174 | No | - | |
| T182 | Yes | Phospho-site | [33] |
| Y187 | Yes | Phospho-site | [4], [5] |
| Y216 | Yes | Phospho-site | [5-10] |
| Y232 | Yes (rat) | Phospho-site | [34] |
| Y233 | No | - | |
| T288 | No | - | |
| Y338 | Yes | - | [11] |
| Y343 | No | - | |
| Y379 | No | - | |
| Y385 | No | - | |
| Y419 | Yes | AutoP | [12-18] |
| Y439 | Yes | Phospho-site | [19-26] |
| Y466 | No | - | |
| Y482 | No | - | |
| Y530 | Yes | Phospho-site | [27-32] |

**Figure S3.** Identification of human c-Src phospho sites in vitro by mass spectrometry as described in experimental procedures

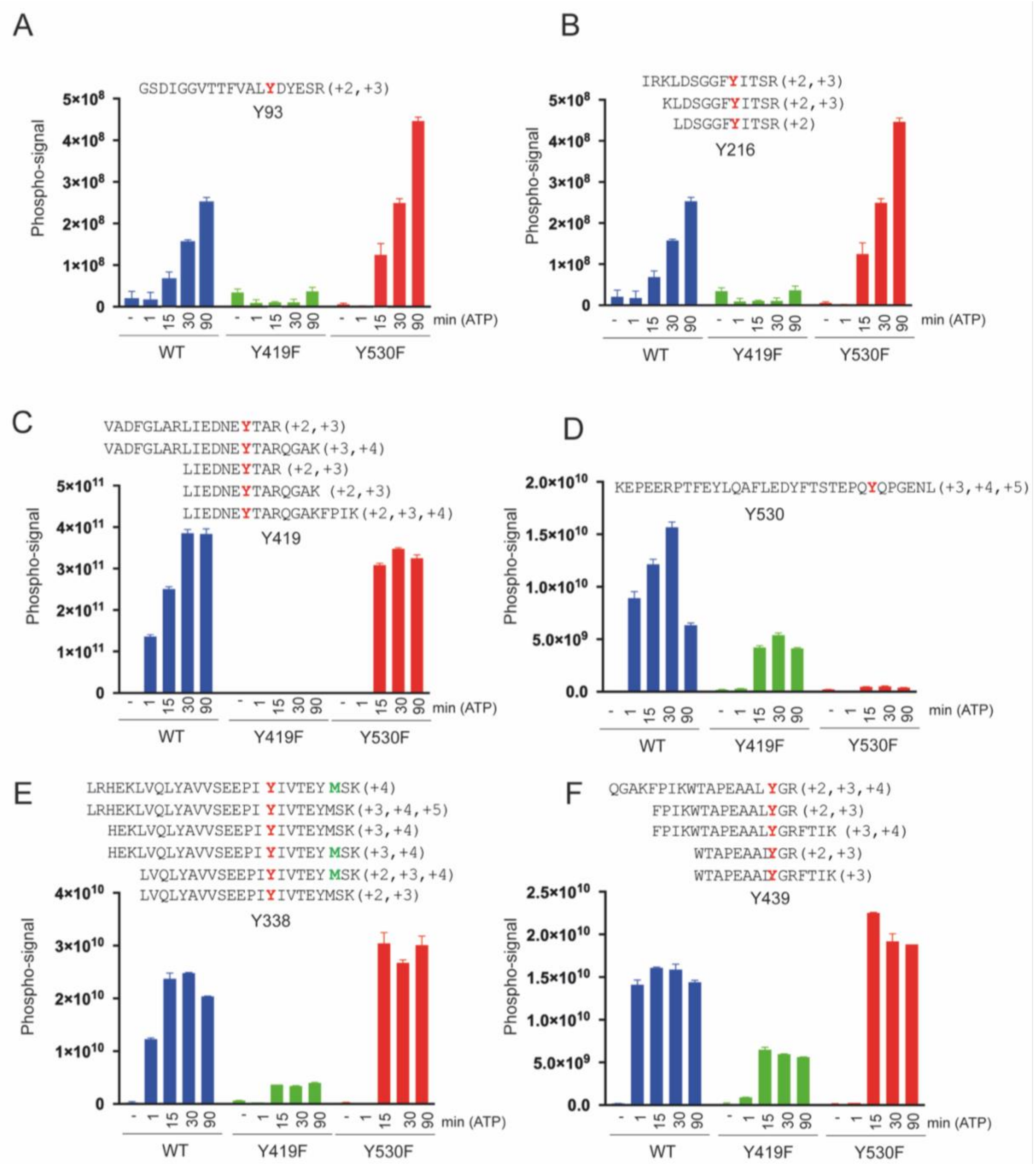


**Figure S4.** Identification of human c-Src phospho sites in vitro by mass spectrometry as described in experimental procedures. Time courses (0-90 min) of c-Src 3D WT, Y419F AND Y530F (2 μM). Normalized phospho-peptide signal (mean ± SD) is representative of 2-3 independent experiments with three technical replicate each. All the phospho-peptide sequences (with their corresponding charge states) used to quantify each phosphorylation site are also shown

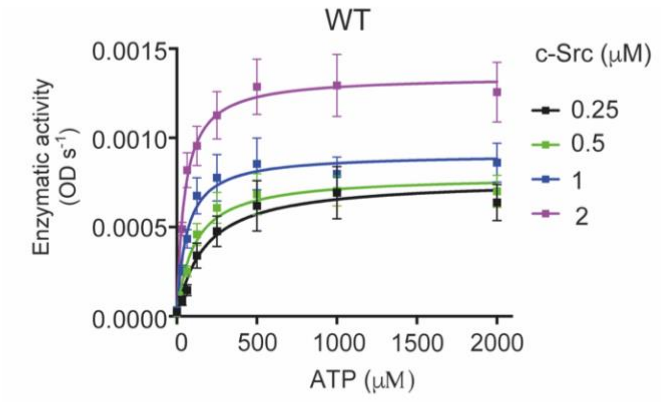


| WT (μM) | 0.25 | 0.5 | 1.0 | 2.0 |
|---|---|---|---|---|
| $V_{max}$ (OD/s) | 0.00076 | 0.00078 | 0.00091 | 0.0013 |
| $K_M$ (μM) | 170.6 | 107.1 | 60.3 | 48.17 |
| $k_{cat}/K_M$ (fold-change) | 1.0 | 1.6 | 3.4 | 6.13 |

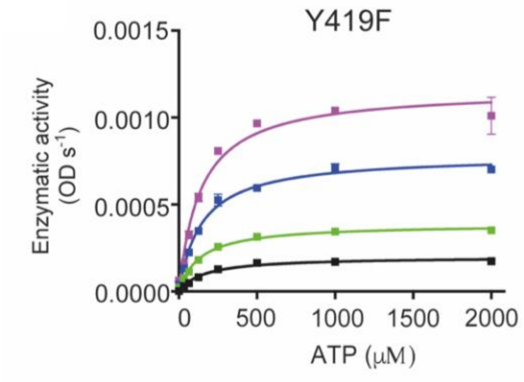


| Y419F (μM) | 0.25 | 0.5 | 1.0 | 2.0 |
|---|---|---|---|---|
| $V_{max}$ (OD/s) | 0.00019 | 0.00038 | 0.00078 | 0.0016 |
| $K_M$ (μM) | 161.5 | 136.1 | 147.0 | 139.7 |
| $k_{cat}/K_M$ (fold-change) | 1.0 | 2.3 | 4.5 | 9.8 |

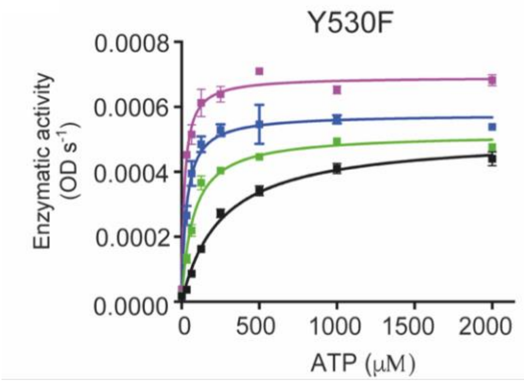


| Y530F (μM) | 0.25 | 0.5 | 1.0 | 2.0 |
|---|---|---|---|---|
| $V_{max}$ (OD/s) | 0.0005 | 0.00051 | 0.00057 | 0.0069 |
| $K_M$ (μM) | 255.4 | 74.3 | 31.3 | 18.3 |
| $k_{cat}/K_M$ (fold-change) | 1.0 | 3.5 | 9.4 | 19.4 |

**Figure S5.** Enzyme kinetics characterization for ATP ($K_M$ ATP) at increasing enzyme concentrations. Enzyme kinetics characterization for ATP at increasing concentrations of enzyme: c-Src (3D) WT, Y419F and Y530F. Table with kinetic parameters and constants with fold-changes (right inset) Data represented are the mean ± SEM of 3-2 experiments in duplicate.

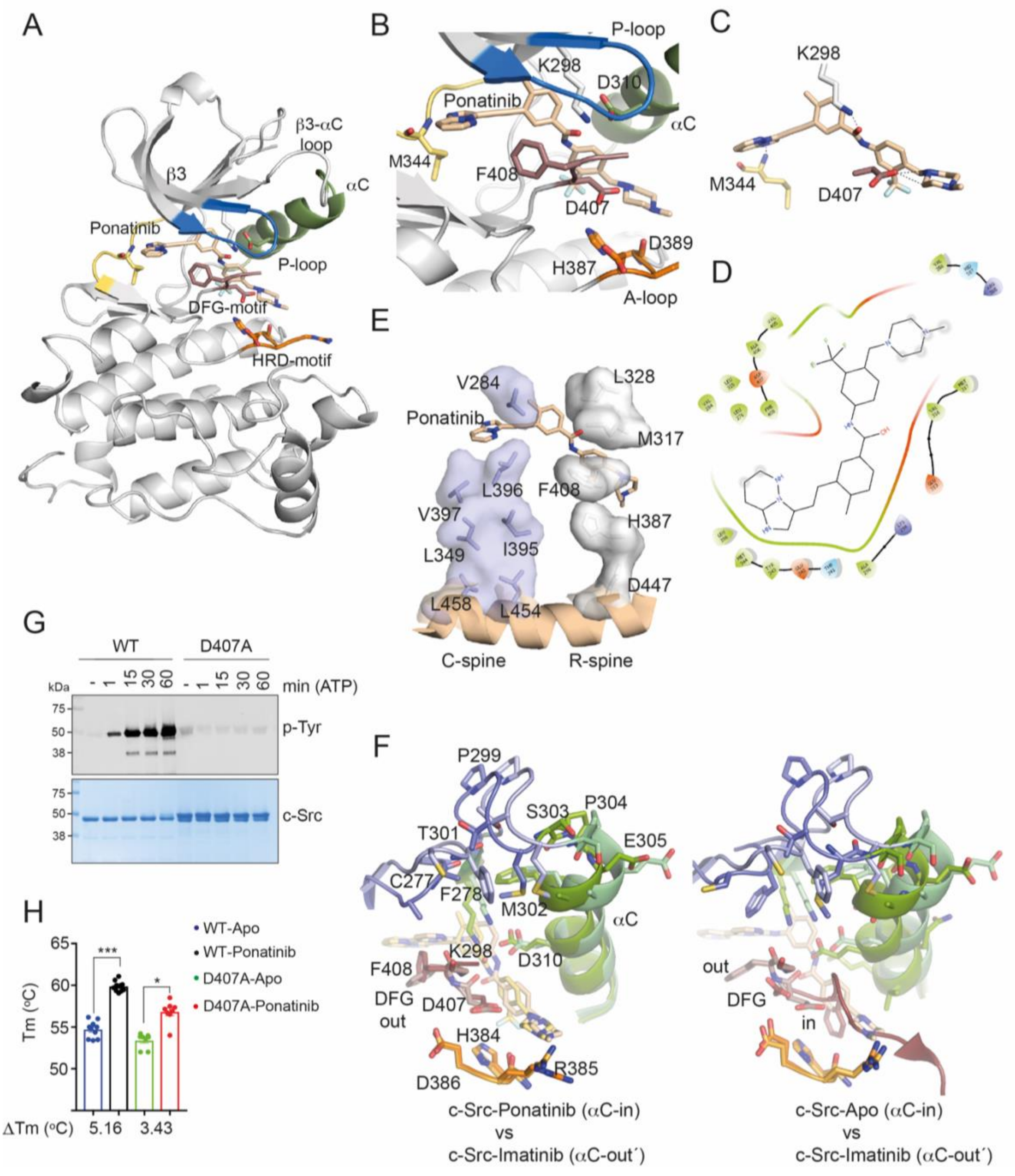


**Figure S6.** Crystal structure of c-Src KD in complex with Ponatinib presents active-like features. (A) Cartoon representation of c-Src KD crystal structure bound to Ponatinib. Selected residues and secondary structure elements are color-coded: P-loop (blue), αC helix (green), hinge (yellow), HRD-motif (orange) and activation segment residues (brown).

(B) Close-up view of the active-site with Ponatinib binding pose showing relevant c-Src interacting residues.

C) Close-up view of Ponatinib binding and c-Src interacting residues.

(D) 2D-pharmacophore representation of Ponatinib.

(E) Cartoon of c-Src catalytic (C-) and regulatory (R-) spines showing side-chain residues and surface representation and how Ponatinib binding breaks the lineal arrangement of the R-spine by flipping F408 from the DFG-motif.

(F) Close-up view of the superimposition of c-Src KD in complex with Ponatinib, c-Src-Imatinib (PDB 2OIQ chain A) and apo (PDB 2OIQ, chain B) crystal structures showing side chain of relevant residues and cartoon representation of secondary structural elements following the same color code as in panel A.

(G) WB of samples from time-course auto-phosphorylation experiment with c-Src WT and D407A (3D-construct 2 $\mu$M) in the presence of ATP (1 mM) and $MgCl_2$ (2 mM) for 0–60 min using a total phospho-tyrosine antibody. Total c-Src protein was evaluated by Coomassie staining.

(H) DSF data of c-Src 3D-construct (WT and D407A) in apo and Ponatinib-complexed states showing the thermal shift (C°). Data represent the mean of the melting temperature (Tm) for each condition ± SEM, out of 4 independent experiment (n= 4) in duplicate. Statistics: ****$p < 0.0001$, one-way test versus control (apo).

**Figure S6 results (continuation)**

Superimposition of our c-Src-Ponatinib crystal structure onto the c-Src-Imatinib complex (Seeliger et al., 2007) revealed some important structural differences at the P-loop, $\alpha$C helix and $\beta$3-$\alpha$C loop (Fig. S6F). Such structural differences could account for the outstanding affinity discrepancies observed between both compounds against c-Src: $K_i$

31 μM and 5.4 nM for Imatinib and Ponatinib, respectively (O'Hare et al., 2009; Seeliger et al., 2007). In the c-Src-Ponatinib crystal structure the P-loop appears to be pointing to the active site contrary to the c-Src-Imatinib complex (PDB code 2OIQ) where it adopts an extended conformation with side chains of F278 and C277 pointing outwards the active site. In this setting M302 from the β3-αC loop is shifted upwards to avoid steric hindrance/clash with F278 and forcing the αC helix to bend adopting an outward configuration (αC-out) and pulling up the whole β3-αC loop in order to avoid clashing with the side chain of M302 (Fig S5F). In the c-Src-Imatinib complex structure (PDB code 2OIQ) the asymmetric unit contained two molecules, where the second protomer lacked the presence of the inhibitor in the active site adopting a DFG-in active configuration, with proper alignment of the R- and catalytic C-spines. Interestingly, this active-like structure displayed the same arrangement of the P-loop, αC and β3-αC loop as the c-Src-ponatinib crystal structure here presented, where F278 (P-loop) side chain is pointing inwards, M302 (β3-αC) is occupying the volume taken by F278 side chain in the imatinib-complexed structure, so there is not steric hindrance for the αC helix to adopt an inner active-like position. (Fig. S6G). It is plausible then, that a thermodynamic penalty caused by an extended P-loop and αC-out configuration account for the low affinity and activity of Imatinib towards c-Src (Seelinger, et al 2007). We validated the effect of mutating to alanine the aspartic acid of the DFG-motif, which formed an important electrostatic interaction with the methyl piperazine group of Ponatinib, in activity and binding assays. As we expected, this mutant precluded significantly the phospho-tyrosine activity of c-Src (Fig. S6H) and had a detrimental binding effect on Ponatinib (Fig. S6I) as indicated by a lower thermal shift measured by DSF.

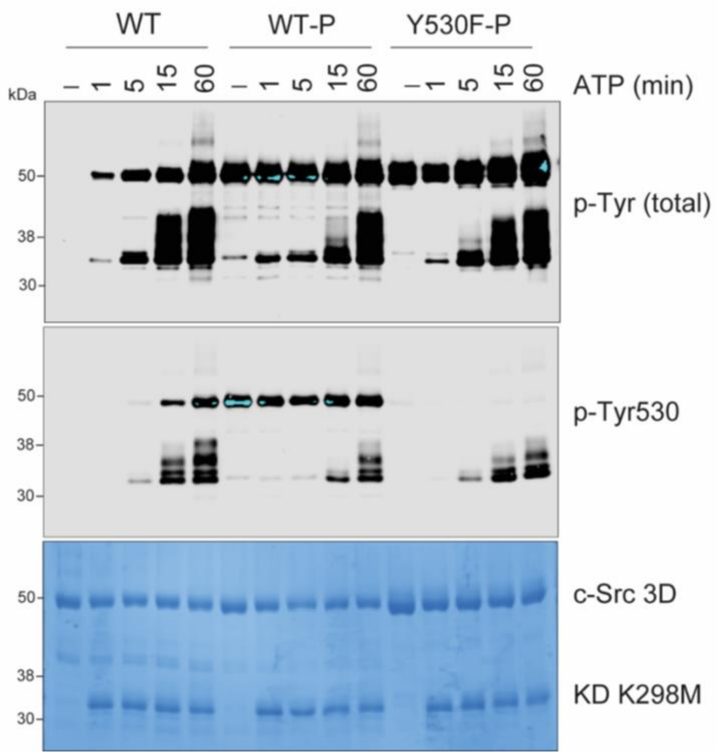


**Fig. S7.** WB analyses of a phosphorylation experiment with c-Src (3D, 1.5 μM) WT and already phosphorylated (90 min) WT and Y530F constructs versus a RET KD K298M (3 μM) as a substrate surrogate using the indicated antibodies. Total protein levels were visualized with Coomassie staining.

### SUPPLEMENTAL TABLES

#### Table 1. Data collection and Refinement Statistics

| | Src KD-ponatinib |
|---|---|
| PDB code | 7OTE |
| Space group | P 1 21 1 |
| Cell dimensions | |
| A, b, c, Å | 42.15, 124.63, 63.59 |
| A, β, γ | 90.00º, 90.15º, 90.00º |
| Resolution (outer resolution shell), Å | 63.59 - 2.5 (2.60-2.5) |
| $R_{sym}$ (%) | 19.0 (94.6) |
| $R_{p.i.m}$ (%) | 8.4 (43.0) |
| I/δ | 7.1 (2.6) |
| Completeness (%) | 93 (100) |
| Redundancy | 4.7 (4.6) |
| No. of unique reflections | 11062 |
| $R_{work}$ | 0.175 (0.189) |
| $R_{free}$[a] | 0.252 (0.256) |
| Total number of atoms | 4388 |
| Wilson B factor | 36.7 |
| Average isotropic B factors, $A^2$ | 44.0 |
| R.M.S.D.S | |
| Bonds, Å | 0.014 |
| Angles, º | 2.066 |
| Ramachandran plot (%) (favored/allowed/disallowed) | 90 / 9 / 1 |

[a] A total of 5.4% of the data were set aside to compute $R_{free}$

**Table S2. SAXs data collection, structural and mass parameter**

| | |
|---|---|
| *Data collection parameters* | |
| Instrument | Diamond Light Source beamline B21 (Harwell Campus, UK) |
| Wavelength (Å) | 1 |
| q-range ($Å^{-1}$) | 0.0032–0.38 |
| Exposure time (s) | 300 |
| Concentration (mg $ml^{-1}$) | 3 |
| Temperature (K) | 293 |
| Structural parameters | |
| Protein | Src-WT |
| Rg (Å) (from Guinier) | 26.12±0.03 |
| Rg (Å) (from P(r)) | 26.10±0.05 |
| Dmax (Å) | 89 |
| *Molecular mass determination* | |
| MM (kDa) from Porod volume | 53 |
| Calculated MM (kDa) from sequence | 50 |
| *Software employed* | |
| Data processing | Scåtter/PRIMUS/ GNOM |
| Ab initio analysis / Averaging | DAMMIF, DAMMIN/DAMAVER |
| Computation of model intensities | FoXS |
| 3D graphics representations | PyMOL |

33. https://www.phosphosite.org/siteAction.action?id=23148314

34.https://www.phosphosite.org/siteGroupAction.action?id=474149&protOrg=661&showAllSites=true&showHTPRefsOnly=true
